# Oncogenic drivers shape the tumor microenvironment in human gliomas

**DOI:** 10.1101/2025.05.16.654515

**Authors:** Bo Zhao, Chae Yun Cho, Lingqun Ye, Tula Keal, Thomas Mitchell, Ines Martin-Barrio, Christina Abi Faraj, Mesut Unal, Jessica Alsing, Nora M. Lawson, Ellie Rahm Kim, Francisco Chavez, Kristy Mendoza Rangel, Sanjay K. Singh, Joseph H. McCarthy, David M. Ashley, W. K. Alfred Yung, Vinay Puduvalli, Guy Nir, Jeffrey S. Weinberg, Linghua Wang, Keith L. Ligon, Giannicola Genovese, Peter Van Loo, P. Andrew Futreal, Jason T. Huse, Jesse R. Dixon, Frederick F. Lang, Kadir C. Akdemir

**Author notes:** Co-first authors (Bo Zhao, Chae Yun Cho, Lingqun Ye, Tula Keal, Thomas Mitchell, Ines Martin-Barrio). Co-senior authors (Jason T. Huse, Jesse R. Dixon, Frederick F. Lang, Kadir C. Akdemir).

## Abstract

Gliomas are aggressive and heterogeneous brain tumors with limited treatment options. While oncogenic mutations in gliomas have been well-characterized, their impact on the tumor microenvironment remains poorly understood. To investigate how genomic alterations may influence the glioma microenvironment, we performed an integrative multiomic and spatial transcriptomic analysis of 93 glioma samples (46 IDH-mutant gliomas and 47 IDH-wildtype glioblastoma) from 69 patients representing both primary and recurrent stages. Using whole-genome sequencing, chromatin conformation capture (Hi-C), RNA-seq, and single-cell spatial transcriptomics (Xenium), we defined how major driver mutations influence spatial tumor organization. We found that IDH-mutant gliomas frequently harbored inflammatory microglia expressing *CX3CR1* specifically within their astrocyte-like malignant neighborhoods. In contrast, glioblastomas demonstrated relatively higher T-cell infiltration and enrichment of immunosuppressive myeloid cell populations. We further compared glioblastomas harboring *EGFR* amplifications due to extrachromosomal DNA (ecDNA) amplifications versus linear chromosomal 7 gains. Tumors with *EGFR* ecDNA displayed increased presence of mesenchymal-like malignant cells, and higher interactions between pericytes and mesenchymal-like malignant cells, likely driven by hypoxia-associated vascular proliferation. Our findings reveal that the mode of oncogene amplification—linear versus ecDNA—shapes distinct tumor architectures and transcriptional dynamics. Taken together, our study highlights the role of oncogenic drivers in shaping the glioma microenvironment, revealing subtype-specific cellular ecosystems that could inform targeted therapeutic strategies.

## Introduction

Gliomas, the most prevalent primary brain tumors in adults, exhibit significant cellular and molecular heterogeneity and clinical outcomes^1^. Current clinical classification of gliomas is based on mutations in the isocitrate dehydrogenase genes (IDH1 and IDH2)^2^. IDH-mutant gliomas, including astrocytomas (IDH-mutant) and oligodendrogliomas (IDH-mut-codel, harboring chromosome 1p/19q deletions), have distinct molecular features and clinical outcomes when compared with IDH-wildtype gliomas, which are predominantly glioblastomas (IDH-wildtype)^3^, emphasizing the importance of incorporating somatic alterations in clinical classification of gliomas.

A hallmark of IDH-mutant gliomas is the accumulation of the oncometabolite D-2-hydroxyglutarate (D-2HG), which leads to widespread DNA hypermethylation phenotype in glioma cells^4^ and consequentially may disrupt molecular processes including chromatin organization around key oncogenes^5–10^ and influence the tumor microenvironment. In contrast, IDH-wildtype gliomas are characterized by higher genomic instability^11^, notably the high frequency of extrachromosomal DNA (ecDNA) amplifications^12,13^. These ecDNA elements drive oncogene heterogeneity and are associated with poorer patient outcomes^14,16^. Recently, several multi-omic analyses of gliomas have deepened our understanding of glioma tumors^17–24^. For example, single-cell RNA sequencing-based studies revealed four primary cellular states: neural progenitor-like (NPC), oligodendrocyte progenitor-like (OPC), astrocyte-like (AC), and mesenchymal-like (MES) enriched in glioma tumors^20^. IDH-mutant gliomas are enriched in OPC-like malignant cell states, whereas IDH-wildtype gliomas predominantly exhibit AC-like and MES-like malignant cell states. The distribution of these cell states influences tumor morphology; for instance, spatial transcriptomics studies of gliomas have demonstrated that tumors with hypoxia-enriched MES-like cell states display more organized compositions compared with their less hypoxic counterparts^21,22^.

Despite these insights, the impact of somatic mutations on tumor composition and spatial architecture remains incompletely understood, particularly in gliomas harboring ecDNAs amplifications which can only be reliably detected through whole-genome sequencing approaches. There is a pressing need for comprehensive multi-omic studies that integrate somatic genome alterations in gliomas with spatial characterization of the tumor microenvironment to elucidate how driver mutations influence the glioma tumor microenvironment. To address this need, we performed a multi-omic analysis of 93 gliomas from 69 patients, including longitudinal samples from 19 patients with multiple time points. Of these 93 gliomas, 47 were primary gliomas, and 46 were recurrent gliomas. Disease recurrence times in this cohort spanned from 2 months to 22 years (Supplementary Table 1). In total, we profiled 449 unique specimens from these 69 patients across our cohort using different molecular assays (Supplementary Table 2). To better characterize structural variants, ecDNA, and complex alterations, half of the samples (n=39) were also analyzed using Oxford Nanopore long-read whole-genome sequencing. To associate somatic genome alterations with *in* vivo tumor architecture, the majority of patients were also profiled with single-cell spatial transcriptomics via the 10x Genomics Xenium platform (n=62 unique specimen from 53 distinct tumors). To further investigate our findings, select tumors were subjected to additional analyses: single-cell analysis (whole-genome sequencing=5; ATAC-seq and/or RNA-seq n=16), bulk chromatin conformation Hi-C (n=20). We used the adjacent tumor sections for the various profiling methods to reduce the effects of tumor heterogeneity in interpretation of our integrative analysis. Tumor purities ranged from 21% to 89% with a mean of 51%, and tumors with more than 25% purities were considered for our analyses. Throughout the manuscript, we integrated these diverse datasets to assess tumor heterogeneity and clonal architecture, providing a comprehensive understanding of the molecular underpinnings of glioma subtypes on tumor microenvironment.

### Characterization of somatic alterations in glioma samples

To investigate the somatic alterations in our glioma samples, we analyzed bulk whole-genome sequencing data from tumor and matched normal (blood) samples. We built a robust data processing pipeline for identifying somatic single-nucleotide variations (SNV), structural variants (SV), copy number alterations (CNA), and extrachromosomal DNA (ecDNA) amplifications (Methods, Supplementary Figure 1). Overall, we observed somatic alterations that are known to be present in specific glioma subtypes^25^. Specifically, chromosome 1p/19q co-deletions were prevalent in IDH-mutant-codel gliomas, while chromosome 7 amplifications and chromosome 10 deletions were common in both IDH-mutant (astrocytoma) and IDH-wildtype gliomas (Figure 1a, Supplementary Table 3). Somatic single-nucleotide hypermutation (tumor mutation burden > 10/mb) was observed in certain temozolomide-treated gliomas, overlapping with earlier findings^26^. Clinically, the recurrence timelines aligned with established patterns^1^, where patients with IDH-wildtype gliomas progressed more rapidly (median=1.1 years), followed by those with IDH-mutant non-codel gliomas (median =3.4), and patients with IDH-mutant-codel gliomas had the longest time to recurrence (median 4.2 years, see Supplementary Figure 2).

**Figure 1.**
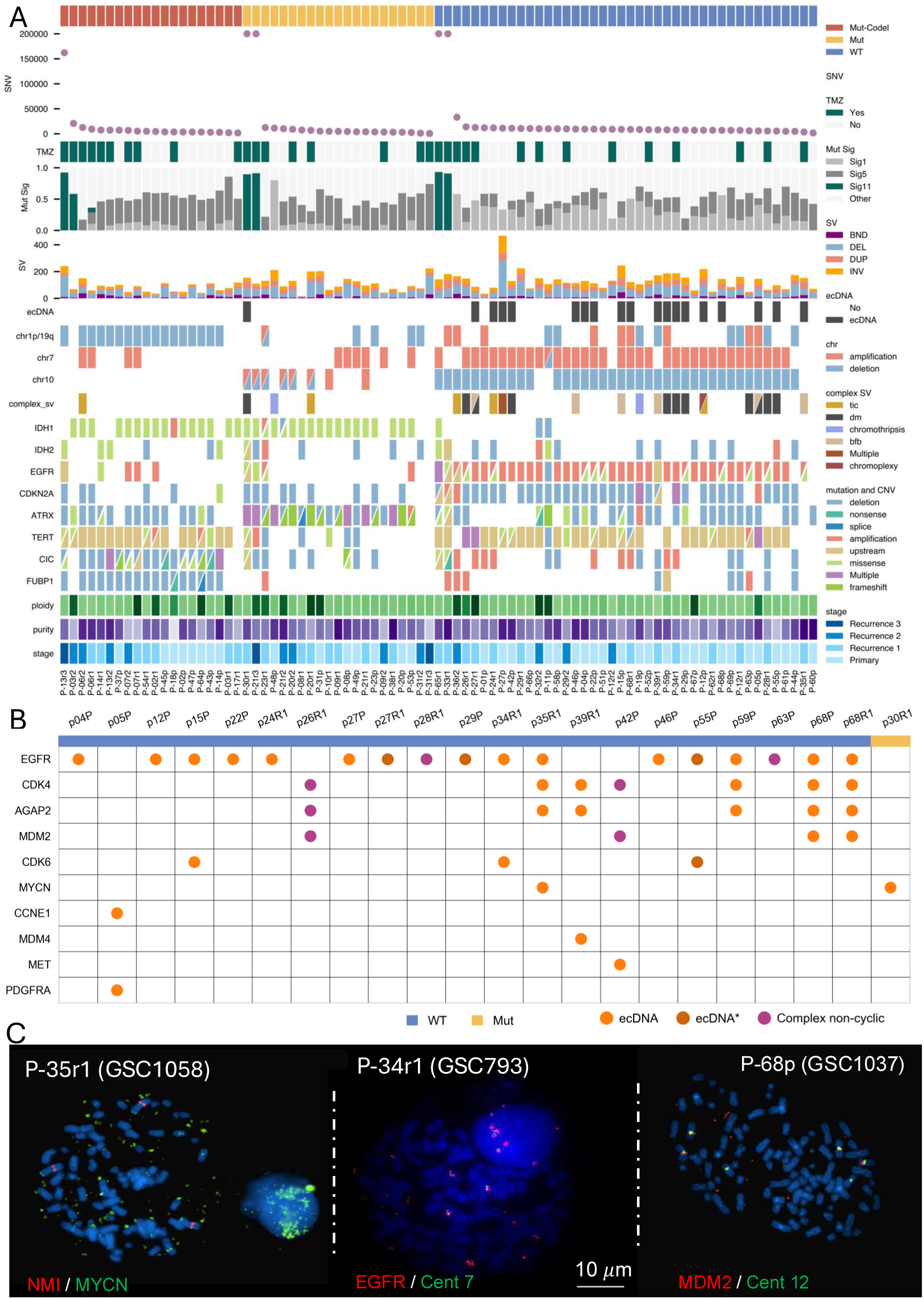
Characterization of somatic mutation landscape in profiled gliomas. **A)** Summary of genomic alterations across glioma samples, stratified by molecular subtype (IDH-mutant-codel, IDH-mutant, IDH-wildtype). Shown from top to bottom are total somatic single nucleotide variants (SNVs), temozolomide (TMZ) treatment status, mutational signature contributions (Signature 1, 5, and 11), structural variant (SV) burden, extrachromosomal DNA (ecDNA) amplification presence, chromosomal-level copy number alterations (chr1p/19q, chr7, chr10), complex SV types, and gene-level mutations and CNAs (e.g., IDH1, EGFR, CDKN2A, TERT). Clinical features including ploidy, purity, and tumor stage (primary, recurrence 1–3) are shown in the bottom heatmaps. **B)** Oncogene amplifications detected by AmpliconArchitect and AmpliconClassifier across glioma samples. Orange circles indicate ecDNA classifications; purple circles denote complex, non-cyclic amplifications lacking evidence for ecDNA topology. * denotes ecDNAs missed by the AmpliconArchitect algorithm but identified using long-read whole-genome and chromatin conformation datasets. **C)** Fluorescence in situ hybridization (FISH) in metaphase spreads of glioma stem cell lines derived from profiled tumors confirms extrachromosomal amplification.

We next examined structural variations (SVs) across our glioma samples and found comparable SV levels among the different subtypes (Figure 1a). Notably, IDH-wildtype gliomas exhibited more complex SVs, characterized by multiple breakpoints within confined genomic regions (Methods). Extra-chromosomal DNAs are frequently observed in glioblastoma genomes but given the complexity of these alterations, whole-genome sequencing approaches are necessary to reconstruct the architecture of such alterations. To characterize complex rearrangements and ecDNAs, we used two alternative methods based on short read sequencing. First, using the AmpliconArchitect (AA) algorithm followed by AmpliconClassifier workflow^27,28^, we identified focal gene amplifications and detected the presence of extrachromosomal DNA (ecDNA) in 15 tumors—14 from IDH-wildtype gliomas and one was a recurrent IDH-mutant tumor—consistent with previous reports on ecDNA prevelance^14–16^ (Figure 1b). Additionally, several tumors exhibited high-level focal amplifications, but AmpliconArchitect did not find sufficient evidence to classify them as circular ecDNA. Second, we also used a graph-based structural variation algorithm, JabBA^29^, to identify potentially missed ecDNA samples by the AmpliconArchtitect algorithm. Three of such tumors were labeled as ecDNA* given the evidence provided by the JabBA algorithm and long-read whole-genome sequencing datasets. Three additional tumors were labeled as complex non-cyclic amplifications (Figure 1b, purple). As a result, 18 tumors were identified with putative ecDNAs in our cohort where the predicted ecDNAs predominantly harbored the *EGFR* oncogene (n=15). In four of these EGFR-amplified tumors, *MDM2* and/or *CDK4* genes were predicted to be co-amplified, while CDK6 was co-amplified in three tumors. Other amplified oncogenes -*CCNE1*, *MET*, *MDM4*, *MYCN*, *PDGFRA*- were found independent of *EGFR*. The single IDH-mutant glioma (p30R1) with ecDNA harbored *MYCN* amplification. This tumor, which had recurred and had been treated with temozolomide, also exhibited SNV hypermutation (226,689 somatic mutations, 75 mutations/megabase), suggesting multiple evolutionary bottlenecks triggered by treatment during disease progression, an observation consistent with previous studies of recurrent IDH-mutant gliomas^15^.

To validate our ecDNA predictions, we utilized patient-derived glioma stem cell (GSC) lines, successfully generated from the some of our profiled tumors. In matched GSCs, we used fluorescence *in situ* hybridization (FISH) to confirm^13^ the high copy number and extrachromosomal nature of detected oncogene amplifications in metaphase spreads (Figure 1c, Supplementary Figure 3).

### Integrative analysis enabled more accurate ecDNA reconstruction

Accurate reconstruction of ecDNA remains challenging in certain tumors, even when using whole-genome sequencing approaches. Given the discrepancies in ecDNA predictions based on short read algorithms, we complemented short-read whole-genome sequencing with high-coverage (>30×) Oxford Nanopore long-read sequencing and bulk Hi-C datasets from ecDNA-predicted tumors. This multimodal dataset enabled a comparative analysis of ecDNA reconstruction algorithms using both short- and long-read whole-genome sequencing in matched tumor samples (Supplementary Table 4).

In tumors with relatively simple ecDNA configurations (e.g., patient P-46p), we observed high concordance between short-read and long-read sequencing reconstructions (Figure 2a). However, tumors with more complex ecDNA structures including those from patients P-59 and P-68 were more difficult to resolve using a single approach. AmpliconArchitect predicted a single ecDNA containing the oncogenes EGFR (from chromosome 7) and CDK4 (from chromosome 12) in both cases. For the P-68 tumor, we observed high concordance between short-read and long-read reconstructions (Figure 2b, Supplementary Figure 4a). In contrast, for P-59, we detected discrepancies between the reconstructions generated by AmpliconArchitect and those from long-read-based algorithms (Decoil^30^ and Coral^31^, Supplementary Figure 4b). While AmpliconArchitect predicted a single circular amplification, long-read sequencing revealed two distinct ecDNA circles—one containing EGFR and the other containing CDK4 (Figure 2c-d). This two-circle model was more consistent with the observed differences in copy number between EGFR (average CN: 91) and CDK4 (average CN: 50) in bulk whole-genome sequencing data. To further evaluate the long-read findings, generated Hi-C data for these samples to characterize their architecture and genomic contiguity. We applied the ec3d algorithm, which integrates Hi-C data for ecDNA reconstruction^32^, to compare the single versus multiple ecDNA configuration predictions. In patient P-59, ec3D analysis showed an increased interaction frequency signal at the corners of the maps reminiscent of circular DNA organization^33^, corresponding to distinct ecDNA circles derived from chromosomes 7 and 12 (Figure 2e-f). In contrast, the AmpliconArchitect prediction of a single circle resulted in an abnormal Hi-C pattern, suggesting that the single-circle reconstruction does not accurately reflect the predominant ecDNA structure in this tumor (Supplementary Figure 4c). Accurate reconstruction of ecDNA is critical for understanding the expression and regulation of ecDNA-residing genes. To investigate enhancer relocation within ecDNAs, we ran neo-loop analysis^34^ using Hi-C data from patient P-59. This showed an enhancer-hijacking event within the chr12-derived ecDNA circle, where *HOXC11* and *HOXC13* genes (chr12: 53.9Mb) were brought into close proximity with a distal enhancer (chr12:20.1-20.2Mb). Both the expression of *HOXC11*, *HOXC13* genes, as well as the accessibility of distal enhancer element, were elevated only in P-59 tumor, compared to other IDH-wildtype gliomas lacking this enhancer-hijacking event (Figure 2g-h).

**Figure 2.**
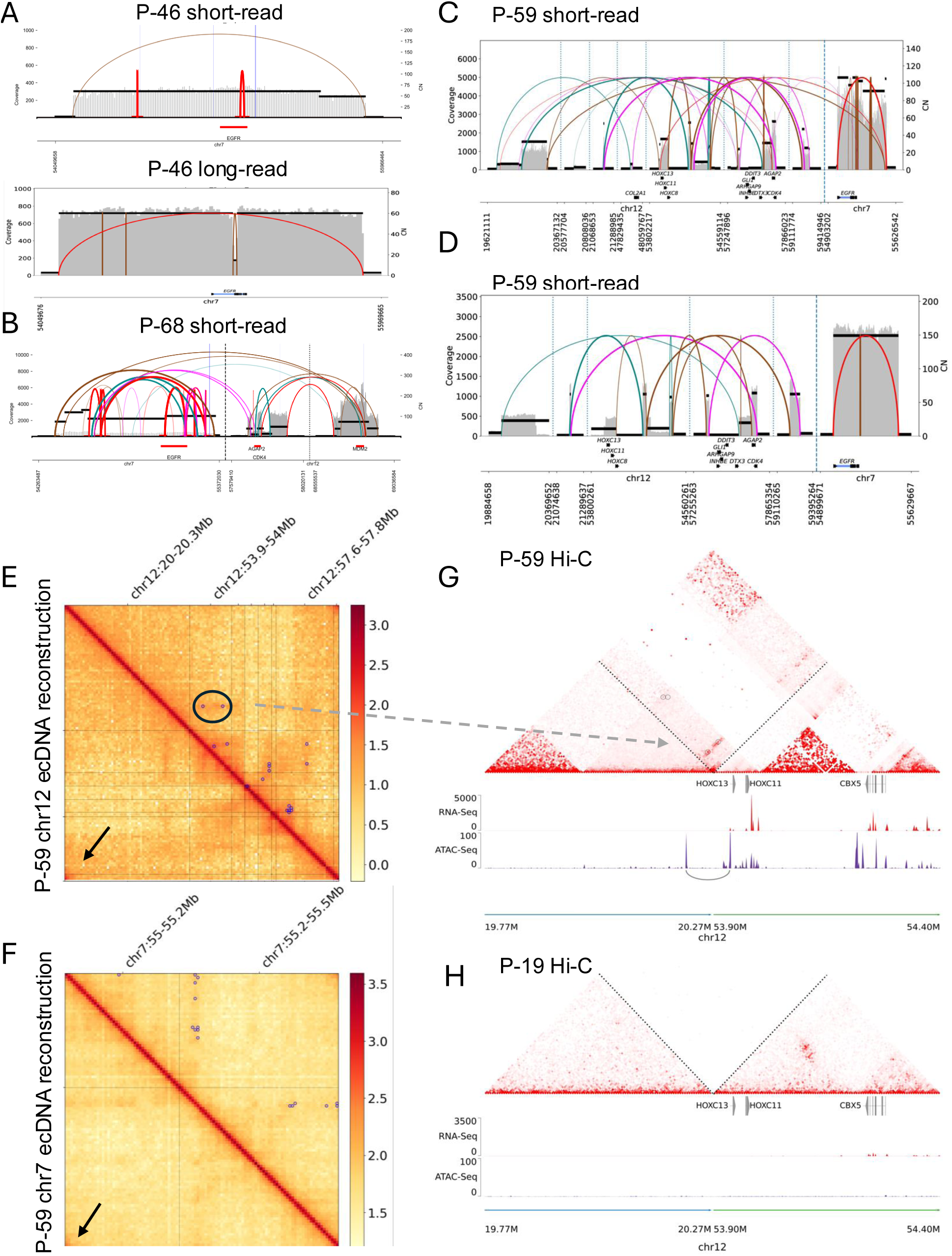
Integrative analysis reveals complex ecDNA architectures and enhancer hijacking event inside an ecDNA in gliomas. **A)** Short-read (top) and long-read (bottom) ecDNA reconstructions for tumor P-46p demonstrate high concordance in a simple EGFR amplification event on ecDNA. **B)** Short-read reconstruction of P-68p reveals a complex ecDNA containing EGFR (chr7), CDK4, and MDM2 (chr12). **C)** Short-read-based prediction for tumor P-59p suggests a single ecDNA circle spanning EGFR (chr7), CDK4, and MDM2 (chr12), but lacks consistency with observed copy number differences. **D)** Long-read sequencing reconstruction of P-59p reveals two distinct ecDNA circles derived from chromosomes 7 and 12, consistent with copy number differences and structural variation. **E–F)** Hi-C interaction maps for P-59p show elevated corner signals (black arrows) indicative of circular architecture in reconstructions of the chr12-derived ecDNA circle (E) and chr7-derived circle (F), supporting the long-read-based ecDNA topology over single-circle prediction. **G)** P-59p Hi-C contact matrix shows neo-loop formation between the HOXC11/13 gene cluster (chr12: ∼54Mb) and a distal enhancer (chr12: ∼20.2Mb), consistent with enhancer hijacking within the chr12 ecDNA. RNA-seq and ATAC-seq tracks show high expression and accessibility at both loci. **H)** A control IDH-wildtype tumor (P-19p) without ecDNA shows no enhancer interaction or activation at these loci.

### ecDNAs contribute to temporal heterogeneity during IDH-wildtype glioma progression

Access to longitudinal samples from the same patients enabled us to study ecDNA dynamics during disease progression. Previous studies have reported that ecDNAs often preserve their structures during progression^16^. Consistent with this, we found that in two patients (P-27 and P-68), both primary and recurrent glioma tumors harbored identical EGFR ecDNAs (Figure 3a, Supplementary Figure 5a-b). Despite radiation and chemotherapy after initial surgery (Supplementary Figure 5c), ecDNA configurations remained unchanged in the recurrent tumors, suggesting limited therapeutic impact on ecDNA-containing clones.

**Figure 3.**
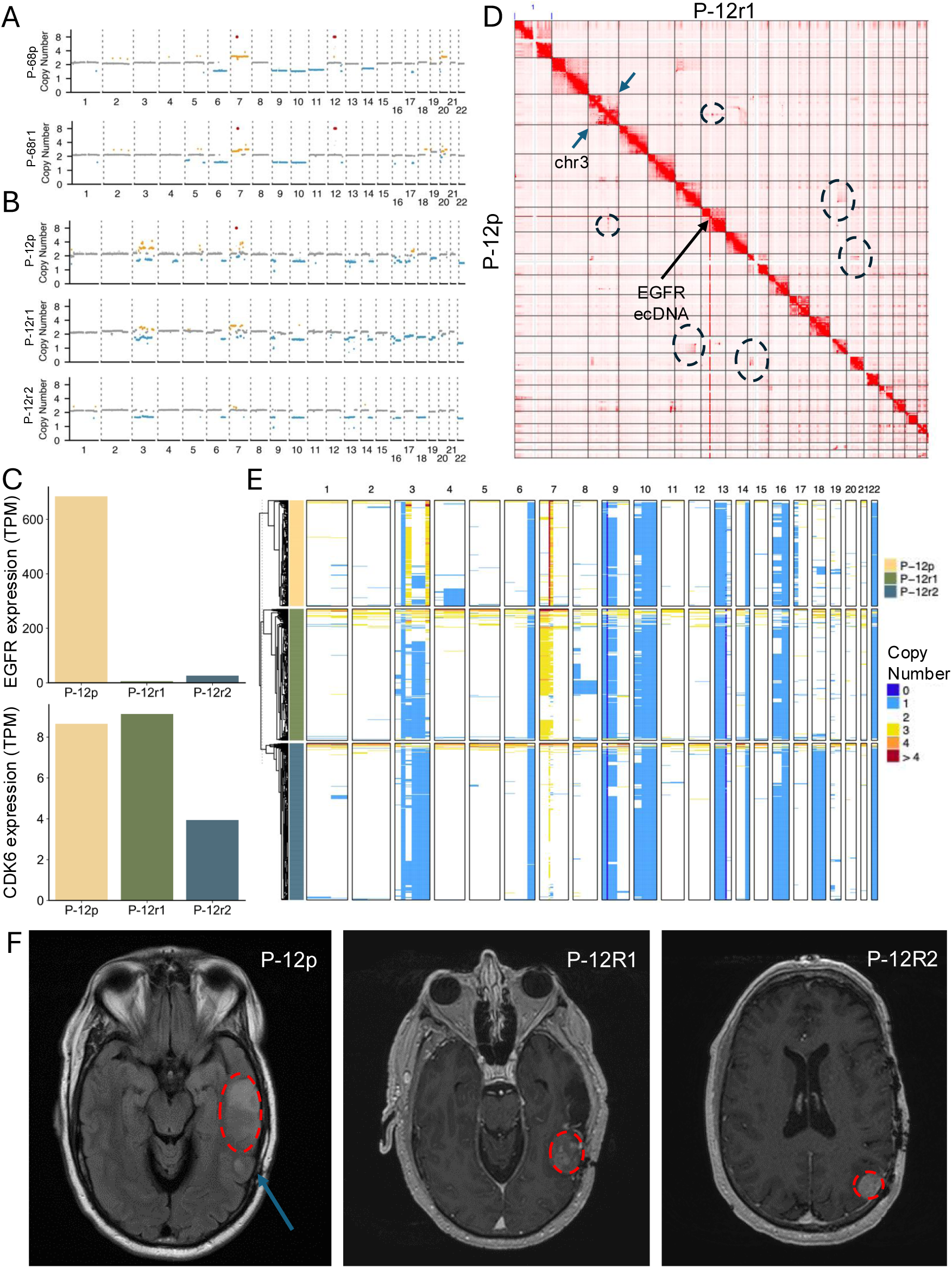
Longitudinal tracking of EGFR ecDNA reveals divergent evolutionary trajectories during glioma progression. **A)** Genome-wide copy number profiles for patient P-68 showing persistence of EGFR ecDNA amplification in both the primary (P-68p) and recurrent (P-68r1) tumors. **B)** Copy number analysis of patient P-12 across three timepoints (P-12p, P-12r1, P-12r2) reveals loss of EGFR ecDNA in recurrent tumors, despite retention of other clonal CNAs. **C)** Bulk RNA-seq analysis shows a marked decrease in EGFR expression in recurrent tumors from P-12, while CDK6 (residing on chromosome 7) expression remains relatively stable, indicating selective loss of the EGFR ecDNA clone. **D)** Hi-C contact matrix from P-12p and P-12r1 shows absence of ecDNA-associated stripe signals in recurrent tumor (upper half matrix) indicated by black arrow. Shared SVs (circles) and chromosome 3 shattering (blue arrows) are indicated for both samples. **E)** Single-nucleus whole-genome sequencing confirms loss of EGFR ecDNA-amplified clones in P-12r1 and P-12r2, with preservation of other chromosomal aberrations (e.g., chr9 and chr10 deletions). **F)** Longitudinal MRIs from P-12 reveal multi-centric disease evolution; recurrent lesions (P-12r1 and P-12r2) emerged outside the initial resection zone (P-12p blue arrow), suggesting spatial heterogeneity and ecDNA clone exclusion from the dominant progression path.

Notably, our longitudinal ecDNA analysis also revealed unusual patterns in two cases. For patients P-12 and P-29, *EGFR* ecDNAs were present in the primary tumors, but absent in the subsequent recurrent tumors, based on copy number profiling, even though other clonal variations were retained (Figure 3b, Supplementary Figure 6a). Since previous studies^16^ reported the emergence of ecDNAs as subclonal events during disease progression or ecDNA persistence in recurrent tumors (as in patients P-27 and P-68), this loss of *EGFR* ecDNA was unexpected. Consistent with the loss of the ecDNA, we observed a marked decrease in *EGFR* gene expression in recurrent tumors from RNA-seq data, while expression of other oncogenes such as *CDK6* also residing on chromosome 7 remained stable (Figure 2c, Supplementary Figure 6b). This suggests that the loss of the ecDNA clone was specific to *EGFR*. To confirm that the lack of ecDNA in the recurrent sample was not due to insufficient sampling of the recurrent samples, we performed multiple additional genomic assays to analyze the presence of ecDNAs in these samples. Hi-C data from these tumors also supported the loss of the ecDNA in the recurrent samples, where the presence of a genome-wide stripe pattern in the primary tumors, a hallmark of ecDNA^35–37^, was absent in recurrent tumors despite sharing other SVs (Figure 3d, Supplementary Figure 6c). Finally, we performed single-nucleus whole-genome sequencing^38^ on primary and recurrent tumor samples from P-12 and P-29, which confirmed the absence of ecDNA-amplified clones in recurrent tumors (Figure 3e, Supplementary Figure 7a,b). However, several other clonal events, such as chromosome 9, 10 deletions in P-12 and chromosomes 6, 10 deletions in P-29 were maintained, indicating that primary and recurrent tumors originated from the same ancestor but diverged following ecDNA loss.

Given the unusual pattern of ecDNA subclone loss in the recurrent tumors, we reviewed the clinical information from these patients. Patient P-12’s first and second recurrences likely originated from an area that was enhancing outside of the primary resection area (Figure 3f, blue arrow). Patient P-29 underwent a partial resection, and recurrences developed in the unresected regions (Supplementary Figure 7c). These findings suggest that the original tumor exhibited spatial and temporal heterogeneity and that ecDNA-containing clones were not the dominant clones leading to disease progression in these cases.

### Glioma genetic drivers are associated with distinct spatial tumor organizations

The observed differences in spatial and temporal ecDNA composition during the evolution of the tumors within our samples highlighted the importance of examining spatial organization of gliomas with different oncogenic driver events. To address this, we performed single-cell spatial transcriptomics (10X Xenium) on matched samples for which we also had whole-genome profiling data. We developed a panel of probes targeting 366 unique genes, enabling the identification of distinct malignant cell states and major cell types within the glioma microenvironment (Supplementary Table 5). In total, we generated over 775 million transcripts from 4.66 million cells across 62 different tumors (Supplementary Table 6). First, we evaluated different cell segmentation methods, including a 5-micron nuclear expansion approach, Baysor^39^, and Proseg^40^ algorithms. Notably, Proseg produced distinct cell state clusters compared with the other methods, offering unique insights into our dataset (Supplementary Figure 8a).

Using our probe set, we were able to characterize major glioma cellular states, including oligodendrocyte progenitor-like (OPC-like), astrocyte-like (AC-like), and mesenchymal-like (MES-like) malignant cell populations. Additionally, non-malignant microenvironmental components such as pericytes, myeloid-derived cells, and T cells were mapped across different glioma subtypes by utilizing a consensus non-negative matrix factorization method (Figure 4a-b). To quantify the spatial organization of these cellular states in different glioma subtypes, we annotated each cell based on the highest program usage (Methods). This analysis showed that IDH-mutant gliomas are enriched in OPC-like cell states compared with IDH-wildtype gliomas, which contain significantly higher MES-like malignant cells, in agreement with previous publications^21–22^. Additionally, we found that myeloid, pericyte and T-cell cell states are significantly higher in IDH-wildtype gliomas compared to IDH-mutant gliomas (Figure 4c). T-cell infiltrations in gliomas has been noted to be lower compared to other solid tumor types^41^, therefore previous publications focusing on immune microenvironment of gliomas generally require a sorting step for enriching immune cell types. Our data suggests that single-cell spatial transcriptomics, compared with spot-based methods, offer higher-resolution characterization of the glioma microenvironment (Figure 4d-e). Relative enrichment of T-cell infiltration in IDH-wildtype gliomas compared to IDH-mutant gliomas can be partially explained by the excessive D2HG levels in IDH-mutant glioma microenvironment which is reported as toxic for T-cells^42^. We also observed significant differences in the myeloid cell populations between IDH-wildtype and IDH-mutant gliomas. Using a classification scheme based on a recent study^43^, we identified two major distinct microglial subtypes in our cells: inflammatory microglia expressing *CX3CR1* and immunosuppressive scavenger cells expressing *CD163* (Supplementary Figure 8b-c). Immunosuppressive microglia were enriched in both IDH-mutant and IDH-mutant codel samples compared to IDH-wildtype gliomas (Figure 4f). In contrast, *CD163*-positive scavenger cells were more abundant in IDH-wildtype tumors, consistent with previous reports (Figure 4f). Interestingly, although IDH-mutant gliomas had a higher proportion of OPC-like malignant cells, microglia cells, particularly the inflammatory microglia, were preferentially enriched within AC-like malignant cell neighborhoods and were depleted in OPC-like neighborhoods within the same tumor specimen, suggesting that the microglial response is shaped by the malignant cell state within the tumor microenvironment (Figure 4g, Supplementary Figure 8d–h). Lastly, to understand the spatial organization of these tumors, we computed a coherence score (Methods), which showed that IDH-wildtype gliomas, harboring higher MES-like cell states, also exhibit more structured tissue organization compared with IDH-mutant, overlapping with previous findings^22^ (Figure 4h-i).

**Figure 4.**
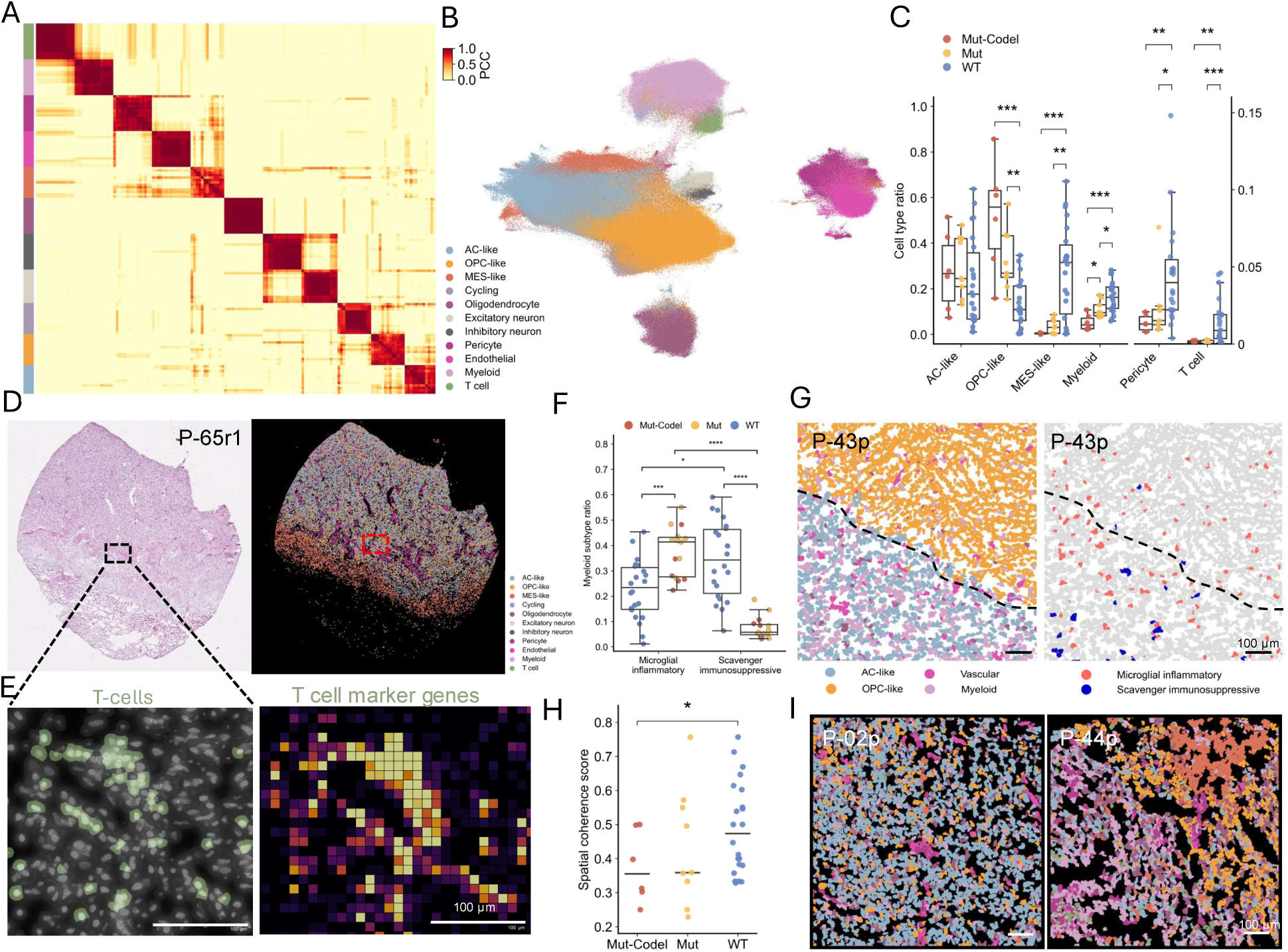
Glioma subtypes have distinct spatial tumor organizations. **A)** Correlation matrix of cell state programs derived from consensus non-negative matrix factorization (cNMF) across spatial transcriptomic profiles highlights distinct malignant and non-malignant states, including OPC-like, AC-like, MES-like, immune, and vascular compartments. **B)** UMAP embedding of over 4 million spatially resolved cells from 62 glioma tumors colored by annotated cell type identities. **C)** Comparison of cell type ratios across IDH-mutant-codel, IDH-mutant non-codel, and IDH-wildtype gliomas shows higher OPC-like states in IDH-mutant and increased MES-like, myeloid, pericyte, and T cell populations in IDH-wildtype tumors. P-values were calculated using the two-sided Wilcoxon rank-sum test (* *P* < 0.05, ** *P* < 0.01, *** *P* < 0.001). **D)** Representative H&E image (left) and Xenium spatial transcriptomic map (right) from an IDH-wildtype (P-65r1) glioma tumor section annotated with major cell types. **E)** High-resolution visualization of T cells (left) and spatial heatmap of T cell marker gene (CCL5, CD2, CD3G, CYTIP, GZMA, PTPRC, TRAC) expression (right) within an IDH-wildype glioma (P-65r1) section. **F)** Myeloid cell population is compared between IDH-wildtype and IDH-mutant (codel and non-codel combined) samples. P-values were calculated using the two-sided Wilcoxon rank-sum test (*** *P* < 0.001). **G)** Representative high-magnification spatial maps from P-43p (IDH-mutant-codel tumor) highlighting the enrichment of myeloid cells within AC-like malignant cell neighborhoods compared to OPC-like malignant cell neighborhood. **H)** Spatial coherence scores quantifying tissue organization show significantly higher values in IDH-wildtype gliomas compared to IDH-mutant subtypes. P-values were calculated using the two-sided Wilcoxon rank-sum test (* *P* < 0.05). **J)** Spatial cell type annotation maps from representative IDH-mutant (P-02p) and IDH-wildtype (P-44p) tumors, illustrating structural differences in tumor organization.

### Extrachromosomal EGFR amplifications lead to distinct spatial tumor organization compared to gliomas with linear EGFR amplifications

Although ecDNAs are frequently observed in IDH-wildtype gliomas, their contribution to spatial tumor organization remains poorly understood. Leveraging the high prevalence of EGFR amplification in our cohort, we utilized our multi-omic framework to stratify IDH-wildtype tumors by the structural context of their *EGFR* amplifications. For this analysis, we focused on the primary tumor’s treatment effects as a confounder. As a result, we identified four tumors with chromosome 7 gains but without ecDNA amplifications of any oncogenes, hereafter referred to as linear amplifications, (e.g., P-67) and five tumors with *EGFR* ecDNAs (e.g., P-46; Figure 5a). Next, we analyzed the cell composition of the microenvironment in these tumors, which showed that tumors with *EGFR* amplifications on ecDNA exhibit significantly higher proportions of MES-like and pericyte cells compared with tumors with linear *EGFR* amplifications (Figure 5b). To validate these results, we also analyzed the bulk RNA-seq data available from the same tumors, which showed that ecDNA-containing tumors have significantly higher hypoxic and metabolic activity compared to linear *EGFR*-amplified tumors (Supplementary Figure 9a). This result differs from the current classification of *EGFR*-amplified gliomas^20,24^, which suggests that these tumors are generally enriched in AC-like cells. While the linear *EGFR* amplification cases do show enrichment of the AC-like cell state, we find that *EGFR* amplifications driven by ecDNA are associated with a high fraction of MES-like cell states. This highlights that our classification of IDH-wildtype gliomas based on whole-genome sequencing provides a more accurate resolution of *EGFR*-amplified subtypes. One key reason for this improvement is that single-cell RNA-seq-based copy number inference methods or exome-sequencing techniques often fail to reliably detect focal, ecDNA-like amplifications. We next examined whether the enriched MES-like and pericyte cells show close spatial proximity in ecDNA-containing tumors (Figure 5c–d, Supplementary Figure 9b). Indeed, in tumors with *EGFR* ecDNA amplifications, MES-like malignant cells and pericyte cells were significantly closer in spatial proximity compared to tumors with linear *EGFR* amplification. The enrichment of pericyte and MES-like malignant cells within shared spatial neighborhoods potentially reflects hypoxia-associated perivascular infiltration. Overall, our results show that ecDNA-containing tumors possess a distinct tumor microenvironment compared to those with linear amplification.

**Figure 5.**
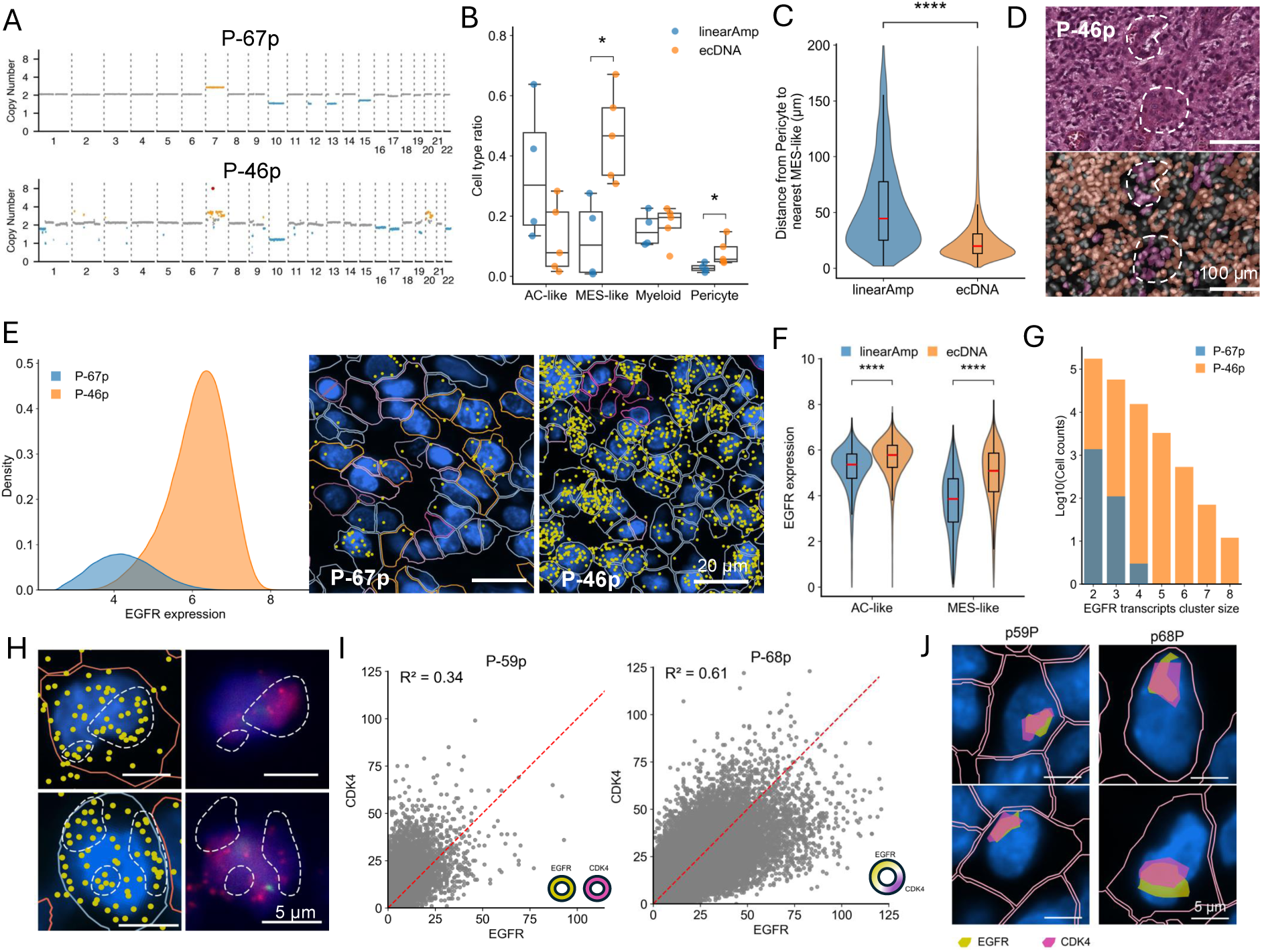
EGFR ecDNA amplifications drives distinct spatial microenvironment and transcriptional clustering in IDH-wildtype gliomas. **A)** Genome-wide copy number profiles of two representative IDH-wildtype gliomas: P-67p with linear amplification of chromosome 7 (top), and P-46p with focal EGFR amplification on ecDNA (bottom). **B)** Cell type composition of tumors with linear versus ecDNA-based EGFR amplification shows significant enrichment of MES-like and pericyte cells in ecDNA cases, while AC-like cells predominate in linear cases. P-values were calculated using the two-sided Wilcoxon rank-sum test (* *P* < 0.05). **C)** Spatial proximity between MES-like and pericyte cells is significantly closer in EGFR ecDNA-containing tumors than in linear amplification tumors. P-values were calculated using the two-sided Wilcoxon rank-sum test (*** *P* < 0.001). **D)** Post-Xenium H&E (top) and spatial transcriptomic cell-type map (bottom) from the same region of P-46p shows MES-like (orange) and pericyte (purple) cells in spatial proximity (highlighted). **E)** EGFR expression distribution is higher in ecDNA (P-46p) versus linear amplified (P-67p) tumors (left). Subcellular mapping (right) reveals significantly higher EGFR transcript levels (yellow dots) in P-46p. **F)** EGFR expression stratified by cell state (AC-like vs. MES-like) shows significantly higher levels in ecDNA tumors compared to linear amplified cases. P-values were calculated using the two-sided Wilcoxon rank-sum test (*** *P* < 0.001). **G)** Distribution of EGFR transcript cluster sizes across cells reveals larger and more frequent transcriptional clusters in ecDNA tumors (P-46p) than in linear amplified tumors (P-67p). **H)** Xenium-detected EGFR transcript clusters (yellow) co-localize with high-copy EGFR regions visualized by post-Xenium FISH from the same cells (magenta) in P-46p, confirming spatial overlap and transcriptional hubs. Dashed white lines shows identified EGFR transcript clusters. **I)** Scatterplots of EGFR and CDK4 expression in tumors P-59p (EGFR and CDK4 on separate ecDNAs) and P-68p (co-amplified on same ecDNA) show stronger expression correlation in P-68p. **J)** Spatial maps of co-localized EGFR (yellow) and CDK4 (darkpink) clustered transcripts polygon in P-59p (top) and P-68p (bottom) show similar levels of transcript clustering despite architectural differences, suggesting hub formation between distinct ecDNA species.

Given the high copy number heterogeneity introduced by ecDNAs, we examined EGFR expression dynamics in tumors with either linear or ecDNA-based amplifications. This analysis revealed that *EGFR* expression was significantly higher in ecDNA-containing tumors across both AC-like and MES-like malignant cell populations, compared to tumors with linear *EGFR* amplification (Figure 5e,f). Next, we leveraged the subcellular resolution of the Xenium platform to identify *EGFR* transcriptional clusters within individual cells. We defined clusters as the presence of at least three *EGFR* transcripts within a 1-micron radius. *EGFR* ecDNA-containing malignant cells exhibited significantly more transcriptional clusters than cells with linear *EGFR* amplification (Figure 5g). As a control for high transcript counts of *EGFR* in ecDNA-containing cells, we identified *FABP7* gene in our panel which showed similarly high expression levels as *EGFR* in P-46 (Supplementary Figure 9c). Performing the same transcriptional cluster analysis showed that the number of *FABP7* transcriptional clusters was lower than that of *EGFR* in the same tumor (Supplementary Figure 9d), suggesting that the observed clustering is not solely due to high expression levels. In addition, the *EGFR* clusters within ecDNA containing tumors showed substantial nuclear overlap, with a median of 70%, suggesting that transcriptional bursts occur within nuclear regions. To further validate this observation, we performed EGFR FISH on the same post-Xenium slides. FISH confirmed the presence of high EGFR copy number clusters that spatially overlapped with transcriptional clusters identified by Xenium (Figure 5h), supporting the hypothesis that nuclear ecDNA hubs drive localized transcriptional activity^37,44^. Finally, we examined how differences in ecDNA architecture influence oncogene co-regulation. Our integrative analysis using long-read and Hi-C data revealed the complex ecDNA structures in tumors P-59 and P-68, both of which harbored amplifications of EGFR and CDK4. In P-59, EGFR and CDK4 were located on separate ecDNA circles, whereas in P-68, both oncogenes were co-amplified on the same ecDNA molecule (Figure 2). As expected, EGFR and *CDK4* expression showed a stronger correlation in P-68 (r^2^ = 0.61) than in P-59 (r^2^ = 0.34) (Figure 5i,j). *MDM2* was amplified only in the ecDNA of P-68, and we observed a strong correlation between *EGFR* and *MDM2* expression in P-68 (r² = 0.81), compared to a much weaker correlation in P-59 (r² = 0.07) (Supplementary Figure 9e). We also checked the clustered transcripts of *EGFR* and *CDK4* in both cases. This data showed despite being on two different circles in P-59, clustered transcripts of *EGFR* and *CDK4* genes are overlapping in similar levels observed in P-68 cells (Figure 5k, Supplementary Figure 9f), suggesting these distinct ecDNAs can form hubs across tumor microenvironment as reported in cancer cells containing different ecDNA species^44^. These results support the accuracy of our ecDNA reconstruction and highlight the importance of ecDNA architecture in modulating oncogene co-expression within the tumor microenvironment.

## Discussion

Our integrative multi-omic and spatial analyses reveal that oncogenic drivers in gliomas exert profound effects on both tumor cell-intrinsic programs and the composition and organization of the tumor microenvironment. In this work, we particularly focused on the effects of IDH mutations and extrachromosomal DNA amplifications in the glioma microenvironment.

IDH mutations lead to excessive accumulation of oncometabolite, D2HG, in the tumor microenvironment. Our analysis showed that *IDH*-mutant gliomas harbor significantly reduced T-cell infiltration compared with *IDH*-wildtype gliomas, although the level of T-cell infiltration is low for wildtype gliomas compared with other solid tumors. Recent work^42^ showed that D2HG can alter T-cell metabolism and reduced T-cell proliferation. Similarly, D2HG has been reported to affect NF-KB signaling and subsequently elevate *CX3CR1* expression in IDH-mutant glioma cells^45^. This may partially explain the increased presence of inflammatory microglia in IDH-mutant glioma microenvironment. Although IDH-mutant tumors were enriched in OPC-like malignant states, inflammatory microglia expressing CX3CR1 were preferentially localized within AC-like regions and were significantly depleted from OPC-like territories within the same tumor sections. This suggests that microglial interactions are not solely dictated by tumor genotype but are also shaped by the local malignant cell state.

We demonstrate that the structural context of oncogene amplification, particularly the amplification of *EGFR* on extrachromosomal DNA, plays a pivotal role in shaping transcriptional dynamics, spatial architecture, and cell–cell interactions within gliomas. ecDNAs have been reported to emerge during disease progression and exhibit spatial heterogeneity in gliomas^46–48^. Our longitudinal analyses further show that ecDNA can be selectively lost during disease progression likely due to spatial heterogeneity, where the ecDNA-containing subclone is resected in the primary tumor, while the remaining non-ecDNA clones drive subsequent recurrences. These divergent trajectories highlight the dynamic nature of ecDNA and underscore the importance of sampling spatially distinct tumor regions to capture ecDNA-related phenotypes. We confirm a high prevalence of *EGFR*-containing ecDNA in these tumors and show that ecDNA amplifications are not only associated with increased copy number heterogeneity but also with spatially organized transcriptional hubs. Using subcellular spatial transcriptomics and combined DNA FISH, we observe that ecDNA-amplified *EGFR* loci form discrete nuclear transcriptional clusters, likely contributing to enhanced gene expression. Importantly, ecDNA-containing tumors are characterized by a microenvironment enriched in MES-like malignant cells and pericytes, as well as increased hypoxic and metabolic activity—features linked to aggressive tumor behavior. Indeed, a recent study which utilized spatial chromatin accessibility technology in a glioblastoma sample also demonstrated that malignant cells with *EGFR* ecDNA molecules are enriched in hypoxic neighborhoods^49^. These findings also overlap with other recent work suggestive of hypoxic features in ecDNA containing pancreatic cancer cells^50,51^. These observations raise a critical question: does hypoxia-induced genomic instability facilitate ecDNA biogenesis, or do ecDNA-containing cells actively remodel the tumor microenvironment to promote hypoxia? Similarly, ecDNA-containing cells may be better adapted to the harsher conditions of hypoxia due to their fitness advantage over cancer cells lacking ecDNA. Understanding the spatio-temporal dynamics of this bidirectional relationship will be essential for understanding ecDNA dynamics and their role in tumor progression.

By integrating short- and long-read sequencing, chromatin conformation capture, and spatial transcriptomic technologies, we provide a comprehensive view of how genomic alterations shape tumor architecture. Our findings highlight that accurate reconstruction^52^ of ecDNA molecules in tumor samples generally requires the combined use of multiple technologies, as each platform, and the algorithms built upon them, carry inherent biases. This integrative approach will be particularly important for identifying and stratifying patients for ecDNA-targeted clinical trials^53^.

Together, our study advances the understanding of glioma biology by revealing how genetic drivers orchestrate distinct cellular ecosystems.

## Acknowledgements

We thank the patients and their families for their selfless contribution to this study. We especially dedicate this work to M. Sachs, whose generosity empowered and further strengthened our commitment to this study. We thank Drs. Sangeeta Goshwami and Preeti Ramadoss for her critical reading of this manuscript. We would also like to thank MD Anderson CATALYST Program and the Brain Tumor Center members, including Lisa Norberg, Chetna Wathoo, Truc Kuo, Stephanie Jenkins, Brittany Parker Kerrigan, Douglas Nielsen, and Jennifer Ritchie, for their essential contributions to patient sample collection and preparation. This work was supported by Brain Cancer SPORE (P50CA127001 to K.C.A., J.T.H., F.F.L), Kleberg Innovative Investigator Award (K.C.A), The Broach Foundation for Brain Cancer Research (to F.F.L), The Elias Family Fund for Brain Tumor Research (to F.F.L), The Priscilla Hiley Cancer Research Fund (to F.F.L), The Sweet Family Brain Cancer Research Fund (to F.F.L), The Jim & Pam Harris Fund, the TLC Foundation From the Heart (to F.F.L), and The Mary Harris Pappas Endowed Fund for Glioblastoma Research (to F.F.L), the Pew Biomedical Scholars Program (to J.R.D.), the Rita Allen Foundation Scholars Program (to J.R.D.), and the 4D Nucleome consortium (U01CA260700 to J.R.D.). This work was also supported by the generous philanthropic contributions to The University of Texas MD Anderson Cancer Center Glioblastoma Moon Shots Program™. We also acknowledge this work utilized services from CPRIT Single-cell CORE Facilities Grant (RP180684) and ATGC core grant (CA016672).

## Data and Code Availability

The raw bulk data reported in this study will be deposited in European Genome Archive. Any additional information is available from the lead contact upon request.

Analysis scripts are deposited in the Akdemir Lab GitHub: https://github.com/akdemirlab/glioma-multiome.

## Author contributions

K.C.A., J.R.D., J.T.H. and, F.F.L. conceived and supervised the study. B.Z. performed the bulk RNA-Seq and spatial transcriptomics analysis and visualization. C.Y.C, T.K, E.R.K, L.Y, and F.C performed the Hi-C data analysis. L.Y. built the somatic mutation analysis pipeline and performed the short-read whole-genome sequencing analysis. T.M. performed the short-read and long-read ecDNA reconstruction analysis. I.M-B. performed the cell line and post-Xenium FISH analysis. K.M.R, C.A.F. conducted clinical annotation. All authors supported clinical contextualization and tumor information integration. J.H.M., D.M.A, K.A.Y, V.P., J.S.W, K.L.L, J.T.H., and F.F.L. provided expert guidance on pathology, and glioma biology. L.W, G.G, P.V.L., and P.A.F. provided critical insights on genomic analysis and interpretation. K.C.A. and J.R.D. wrote the manuscript with input from all authors.

## Supplementary Information

### Supplementary Tables

**Supplementary Table 1**

Clinical characteristics of 69 glioma patients analyzed in this study.

**Supplementary Table 2**

The list of 93 tumor samples collected from 69 glioma patients, comprising 47 primary tumors and 46 recurrent tumors (29 first recurrence, 14 second recurrence, and 3 third recurrence). Samples were classified by IDH status as IDH-wildtype (n = 47), IDH-mutant (n = 25), or IDH-mutant-codel (n = 21). Different data types generated per sample is denoted.

**Supplementary Table 3**

Summary of somatic mutation and structural variant analyses from short-read whole-genome sequencing of glioma samples.

**Supplementary Table 4**

This table compares ecDNA reconstruction results obtained using short-read whole-genome sequencing (AmpliconArchitect) and long-read sequencing analyzed via the Coral algorithm with two support thresholds (≥5 and ≥10 supporting reads).

**Supplementary Table 5**

The list of Xenium gene probe set used in this study.

**Supplementary Table 6**

Overview of single-cell spatial transcriptomic data generated using the 10X Genomics Xenium platform across the glioma cohort.

**Supplementary Figure 1.**
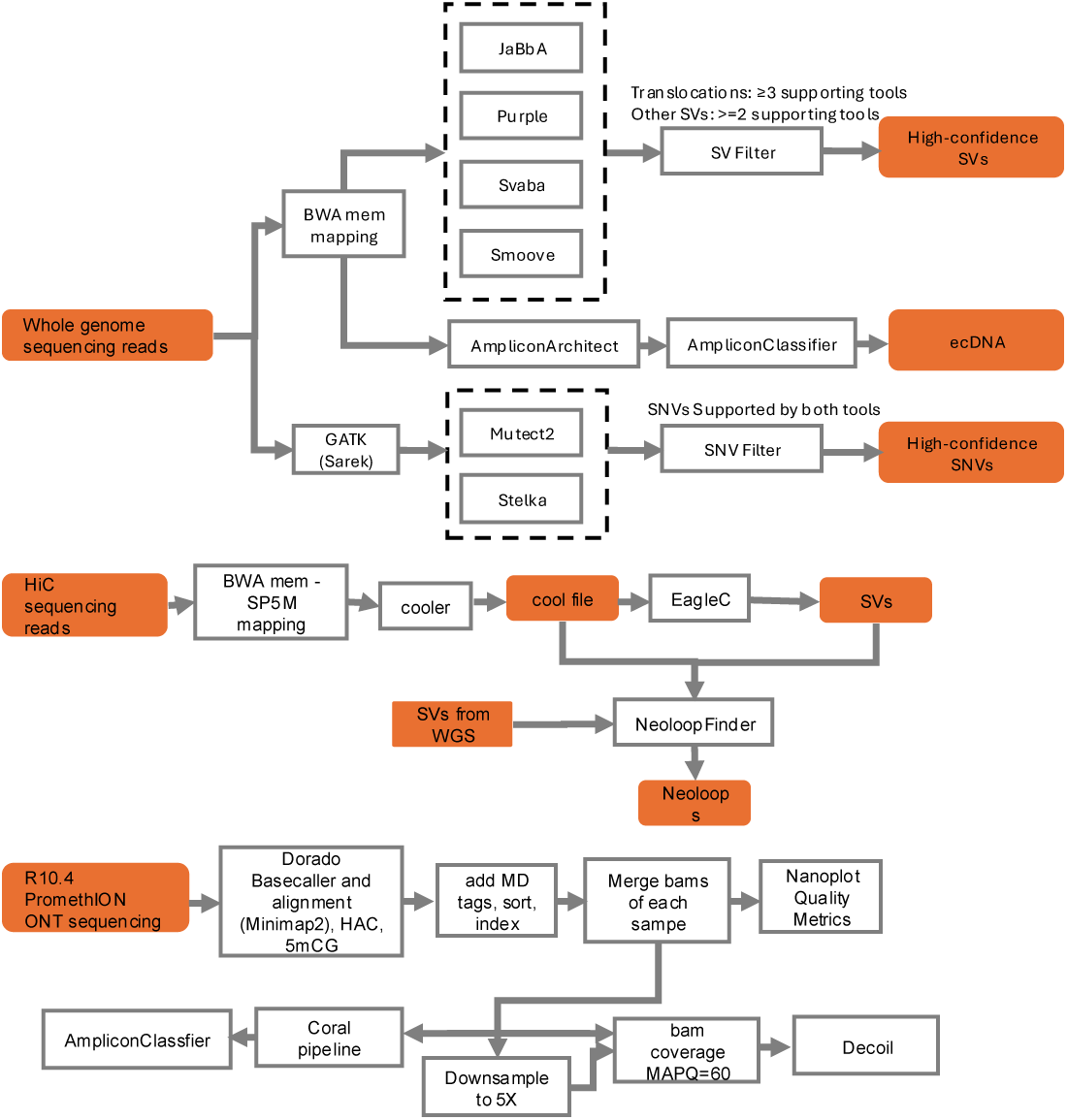
**Overview of the somatic variant and ecDNA analysis pipeline.** This schematic outlines the computational workflow used for identifying high-confidence somatic single-nucleotide variants (SNVs), structural variants (SVs), and extrachromosomal DNA (ecDNA) in glioma whole-genome sequencing data. SNVs were detected using both Mutect2 and Strelka2 within the GATK (Sarek) pipeline, with only concordant variants retained. Structural variants were identified using JaBbA, SvABA, Smoove, and Purple, and filtered based on multi-tool support (≥3 tools for translocations; ≥2 tools for other SV types). ecDNA was reconstructed from high-confidence copy number and breakpoint data using AmpliconArchitect and classified with AmpliconClassifier. This integrative workflow ensured robust detection of somatic alterations and ecDNA across glioma samples.

**Supplementary Figure 2.**
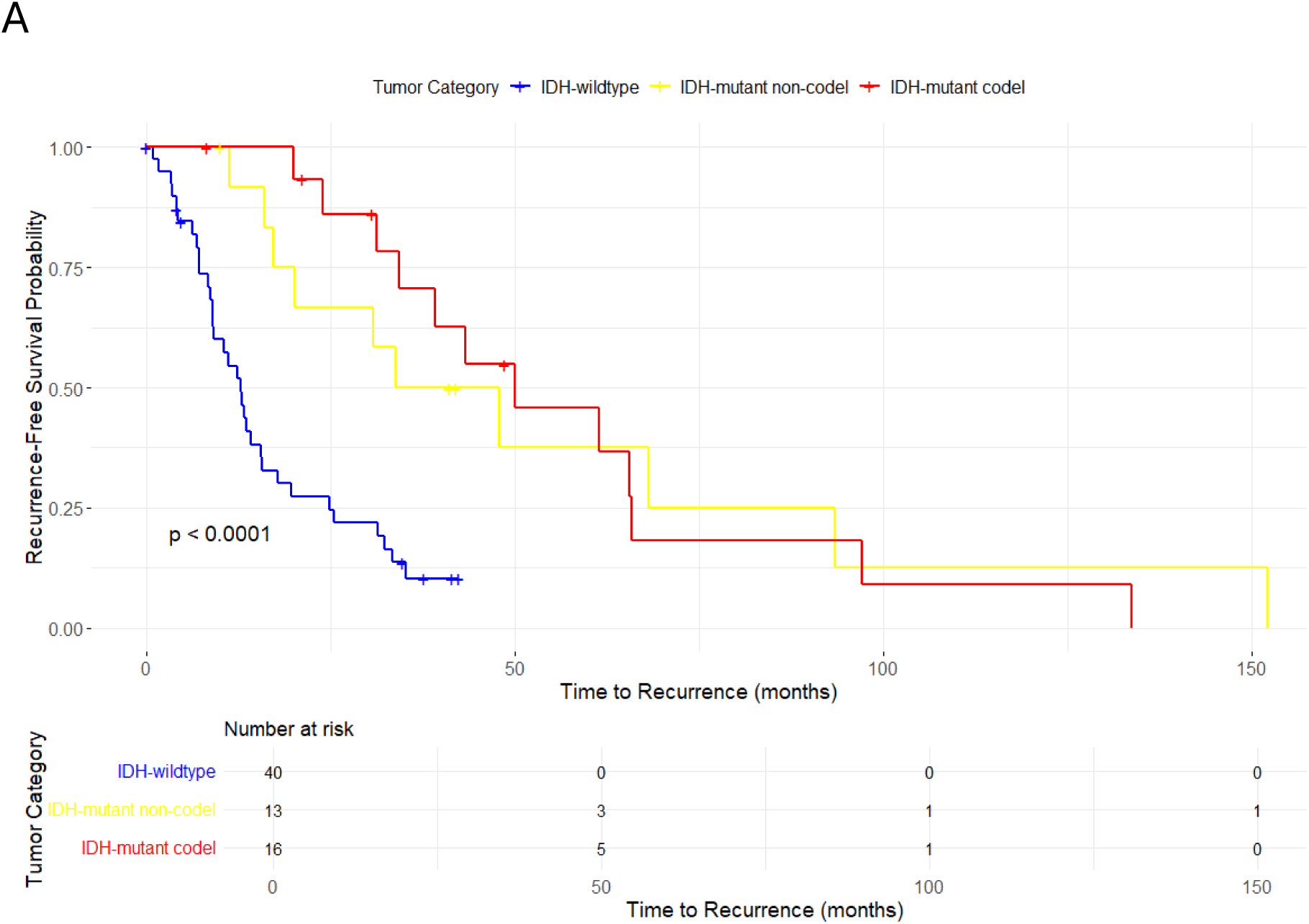
**Survival of patients according to their IDH-mutation classification.**

**Supplementary Figure 3.**
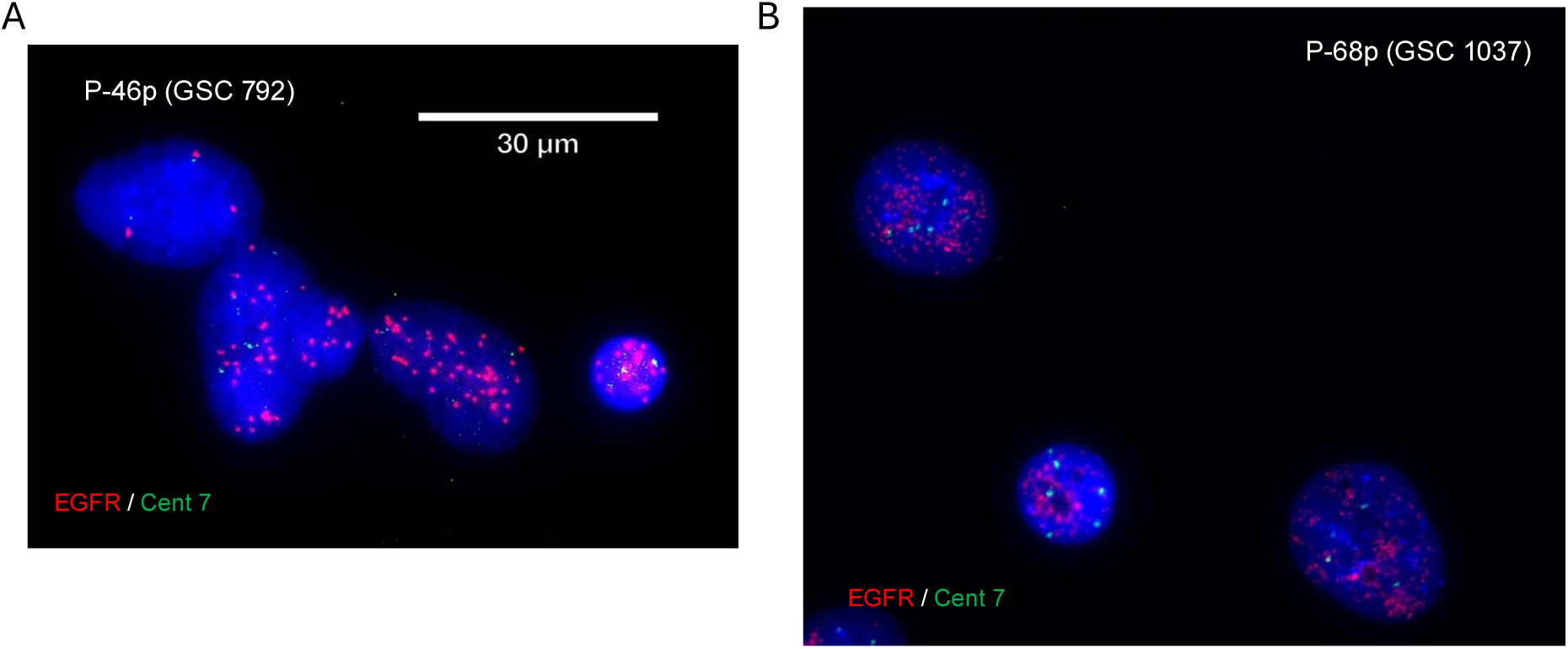
**Fluorescence in situ hybridization (FISH) in interphase cells of glioma stem cell lines derived from profiled tumors showed high oncogene levels reminiscent of extrachromosomal amplification.**

**Supplementary Figure 4.**
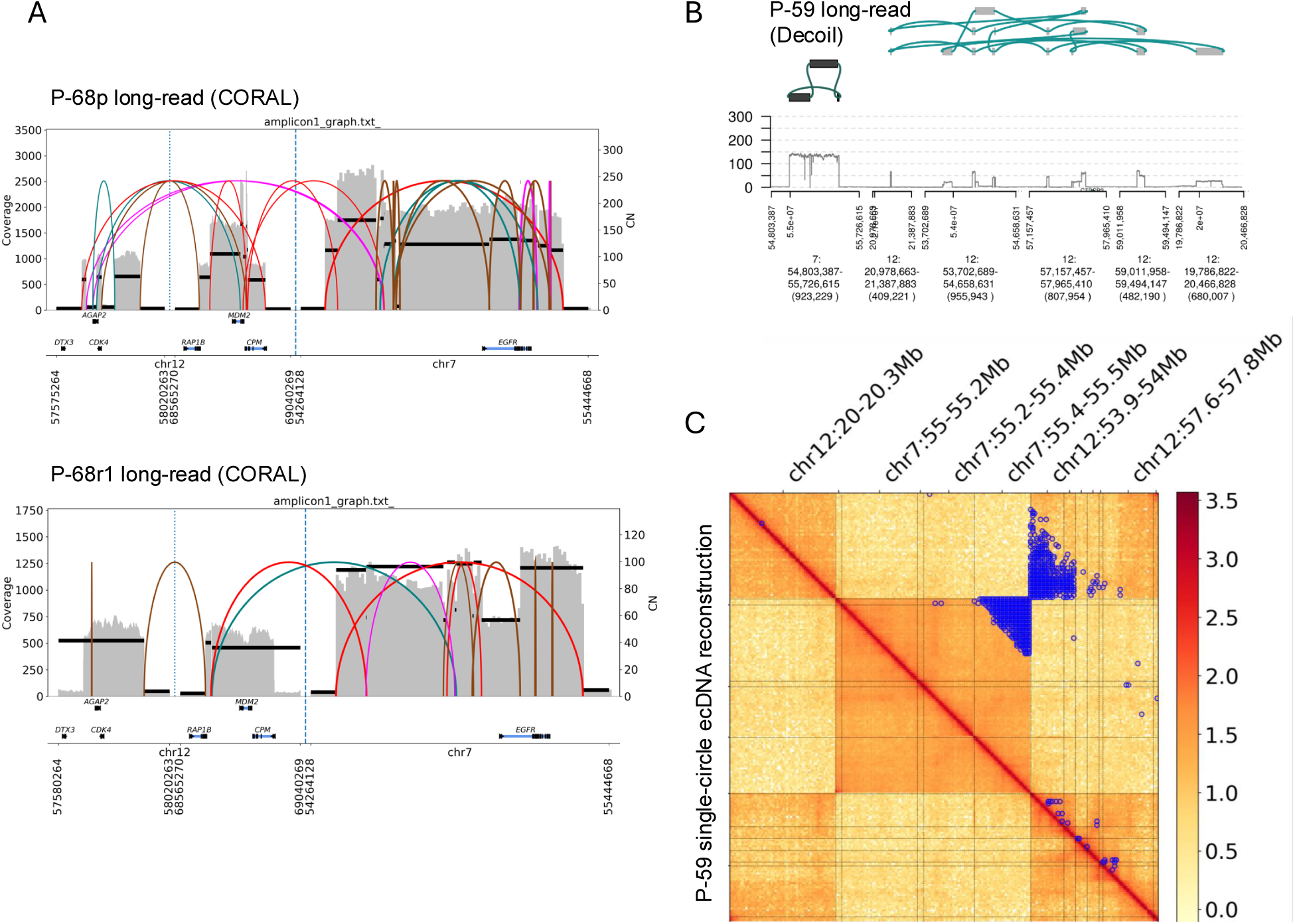
**A)** Long-read ecDNA reconstructions for tumor P-68 primary (top) and recurrent (bottom) demonstrate high concordance in a complex EGFR/CDK4/MDM2 amplification event on ecDNA. **B)** Long-read sequencing reconstruction of P-59p reveals two distinct ecDNA circles derived from chromosomes 7 and 12, consistent with copy number differences and structural variation. **C)** Hi-C interaction maps for the P-59p amplification reconstructed using AmpliconArchitect’s single-circle model showed high variability in off-diagonal signals, suggesting that regions from chromosomes 7 and 12 are unlikely to reside within the same circular structure.

**Supplementary Figure 5.**
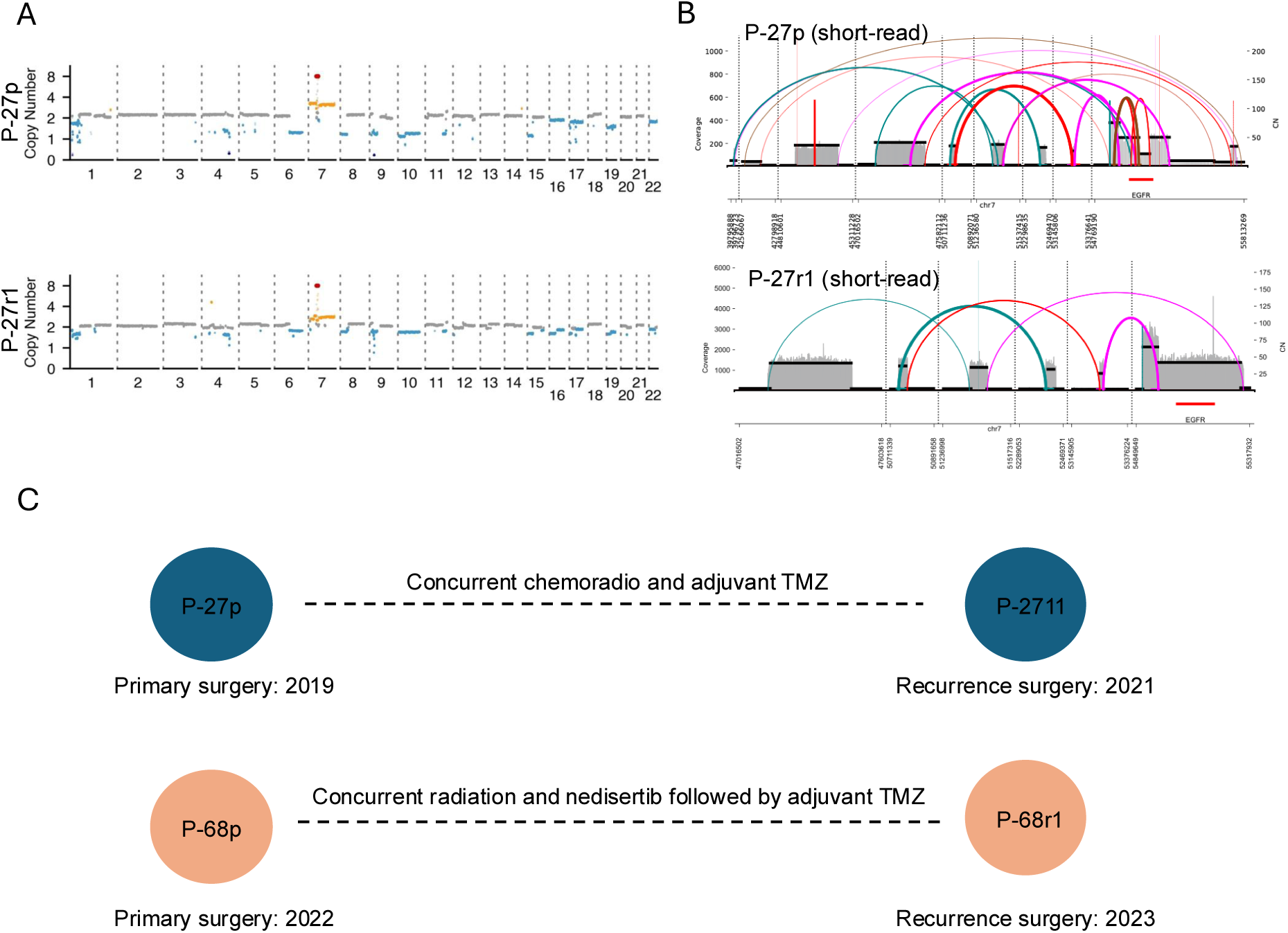
**A)** Genome-wide copy number profiles for patient P-27 showing persistence of EGFR ecDNA amplification in both the primary (P-27p) and recurrent (P-27r1) tumors. **B)** Short-read ecDNA reconstructions for tumor P-27 primary (top) and recurrent (bottom) demonstrate high concordance in a complex EGFR/CDK4/MDM2 amplification event on ecDNA. **C)** Clinical timeline and treatment information for patients 27 and 68.

**Supplementary Figure 6.**
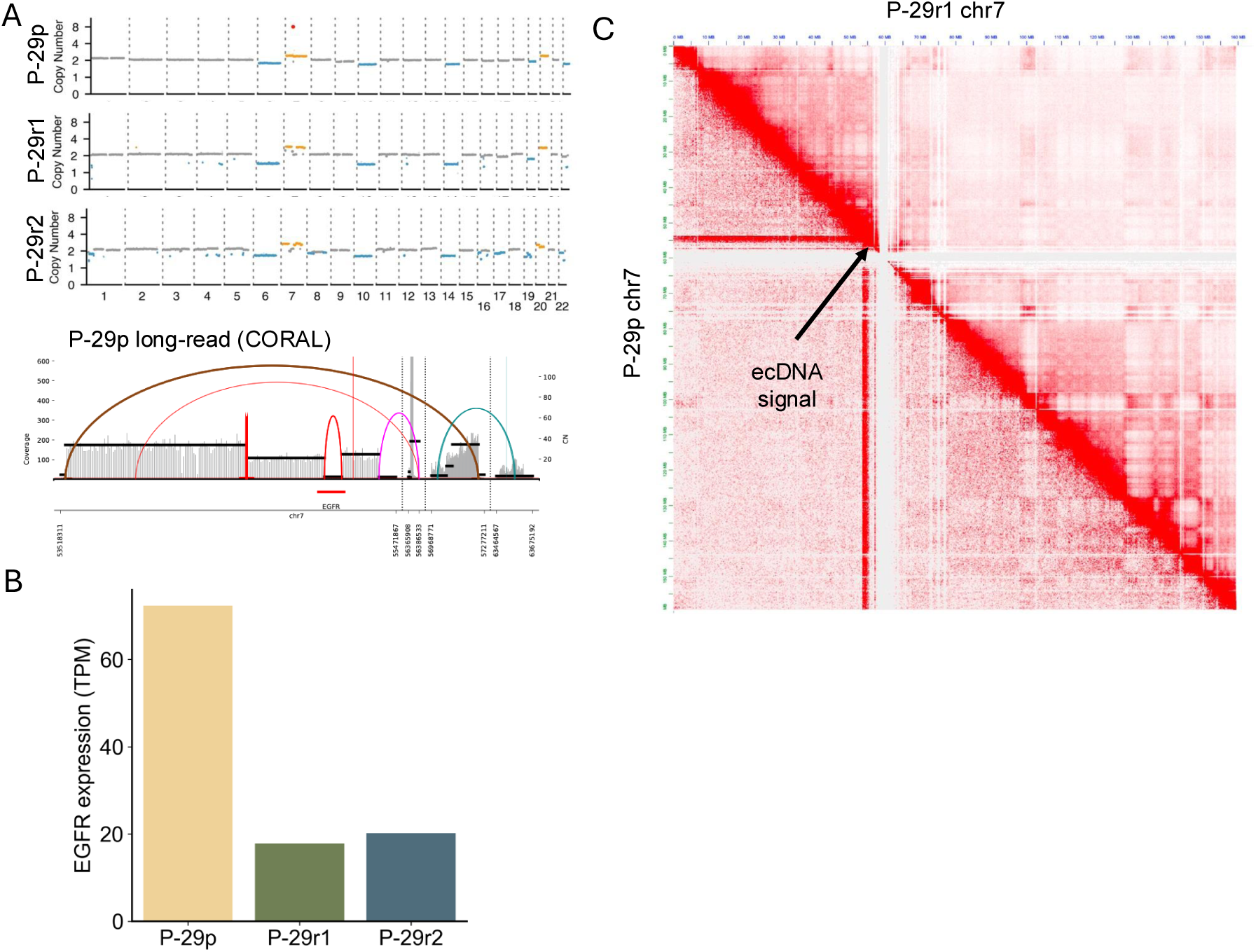
**A)** Copy number analysis of patient P-29 across three timepoints (P-29p, P-29r1, P-29r2) reveals loss of EGFR ecDNA in recurrent tumors, despite retention of other clonal CNAs. Bottom) Long-read sequencing reconstruction of P-29p reveals circular ecDNA structure. **B)** Bulk RNA-seq analysis shows a marked decrease in EGFR expression in recurrent tumors from P-29. **C)** Hi-C contact matrix from P-29p and P-29r1 shows absence of ecDNA-associated stripe signals in recurrent tumor (upper half matrix) indicated by black arrow.

**Supplementary Figure 7.**
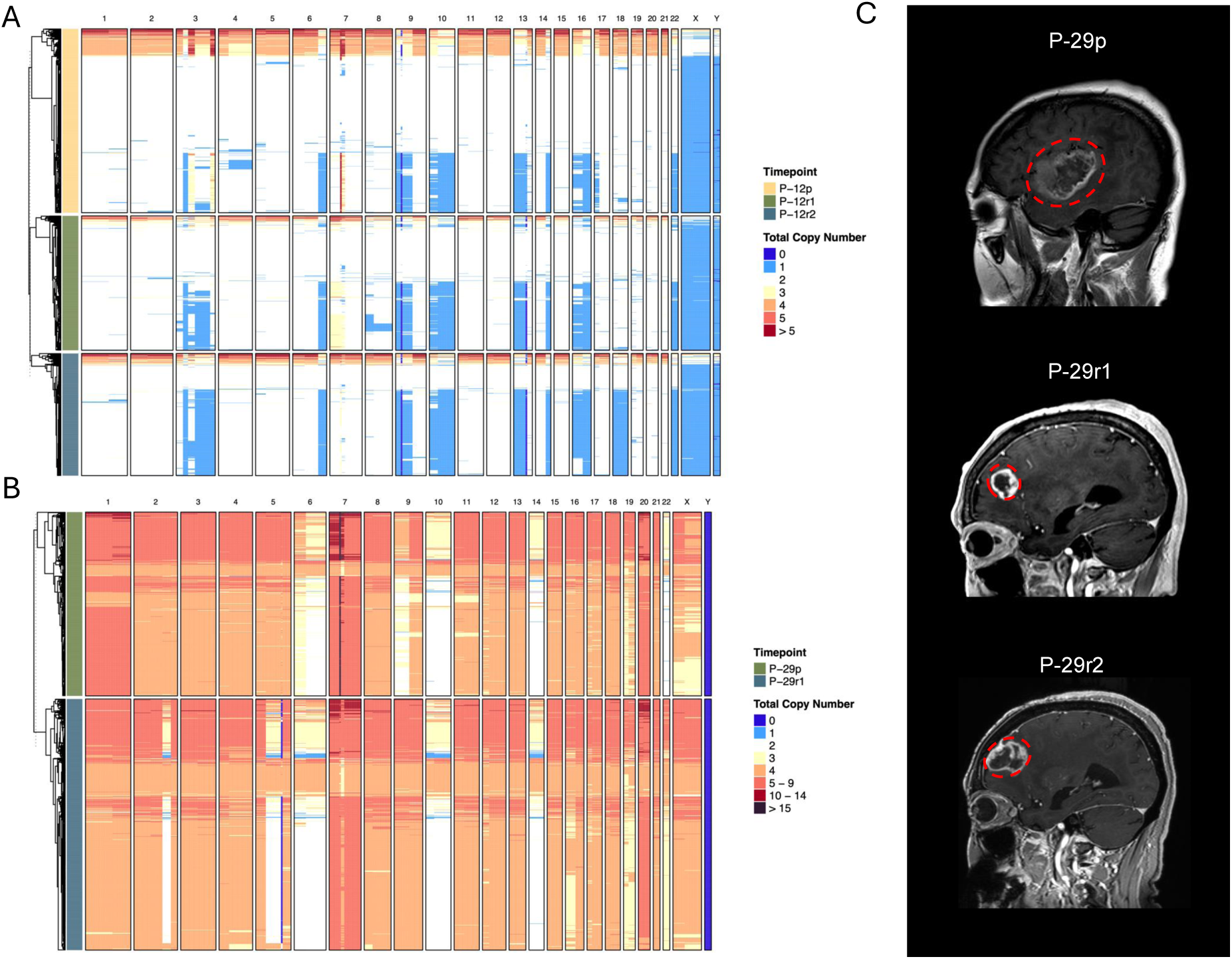
**A)** Single-nucleus whole-genome sequencing confirms loss of EGFR ecDNA-amplified clones in P-12r1 and P-12r2, with preservation of other chromosomal aberrations (e.g., chr9 and chr10 deletions). **B)** Single-nucleus whole-genome sequencing confirms loss of EGFR ecDNA-amplified clones in P-29r1, with preservation of other chromosomal aberrations (e.g., chr6 and chr10 deletions). **C)** Longitudinal MRIs from P-29 reveal multi-centric disease evolution; recurrent lesions (P-29r1 and P-29r2) emerged from the residual tumor area after P-29p surgery, suggesting spatial heterogeneity and ecDNA clone exclusion from the dominant progression path.

**Supplementary Figure 8.**
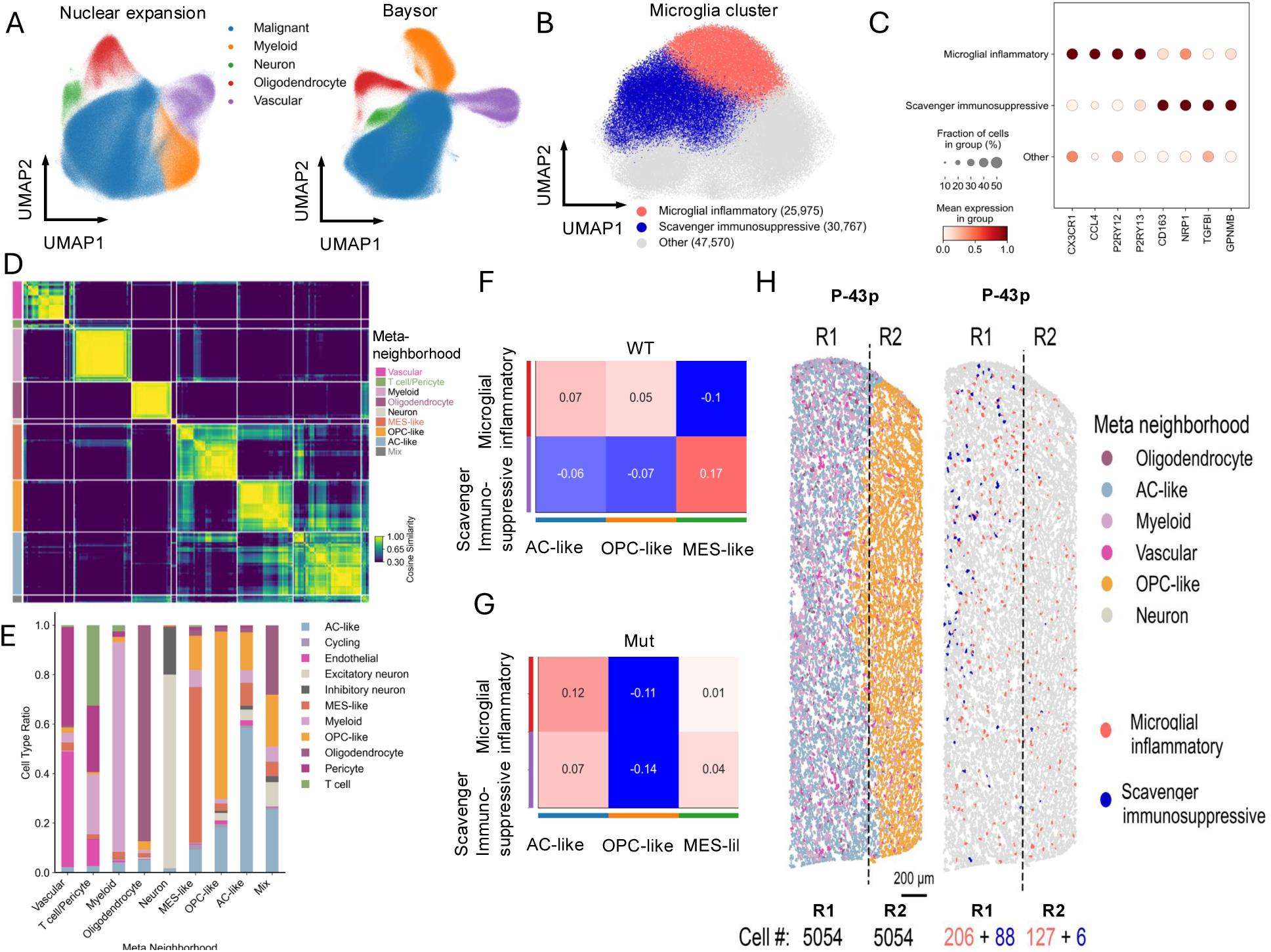
**A)** Comparison of cell segmentation strategies using nuclear expansion, Baysor, and Proseg algorithms. **B-C)** Classification of microglial subtypes based on dominant marker gene expression. **D-E)** Neighborhood analysis to investigate microglia-enriched spatial neighborhoods. **F-G)** Analysis of microglial subtype enrichment within malignant cell neighborhoods. Scavenger immunosuppressive microglia were enriched in MES-like neighborhoods in IDH-wildtype gliomas (F), whereas inflammatory microglia were enriched in AC-like neighborhoods and depleted in OPC-like neighborhoods in IDH-mutant gliomas (G). **H)** Representative spatial map from P-43p (an IDH-mutant-codel tumor) highlighting the enrichment of myeloid cells within AC-like malignant neighborhoods (R1) compared to OPC-like malignant neighborhoods (R2).

**Supplementary Figure 9.**
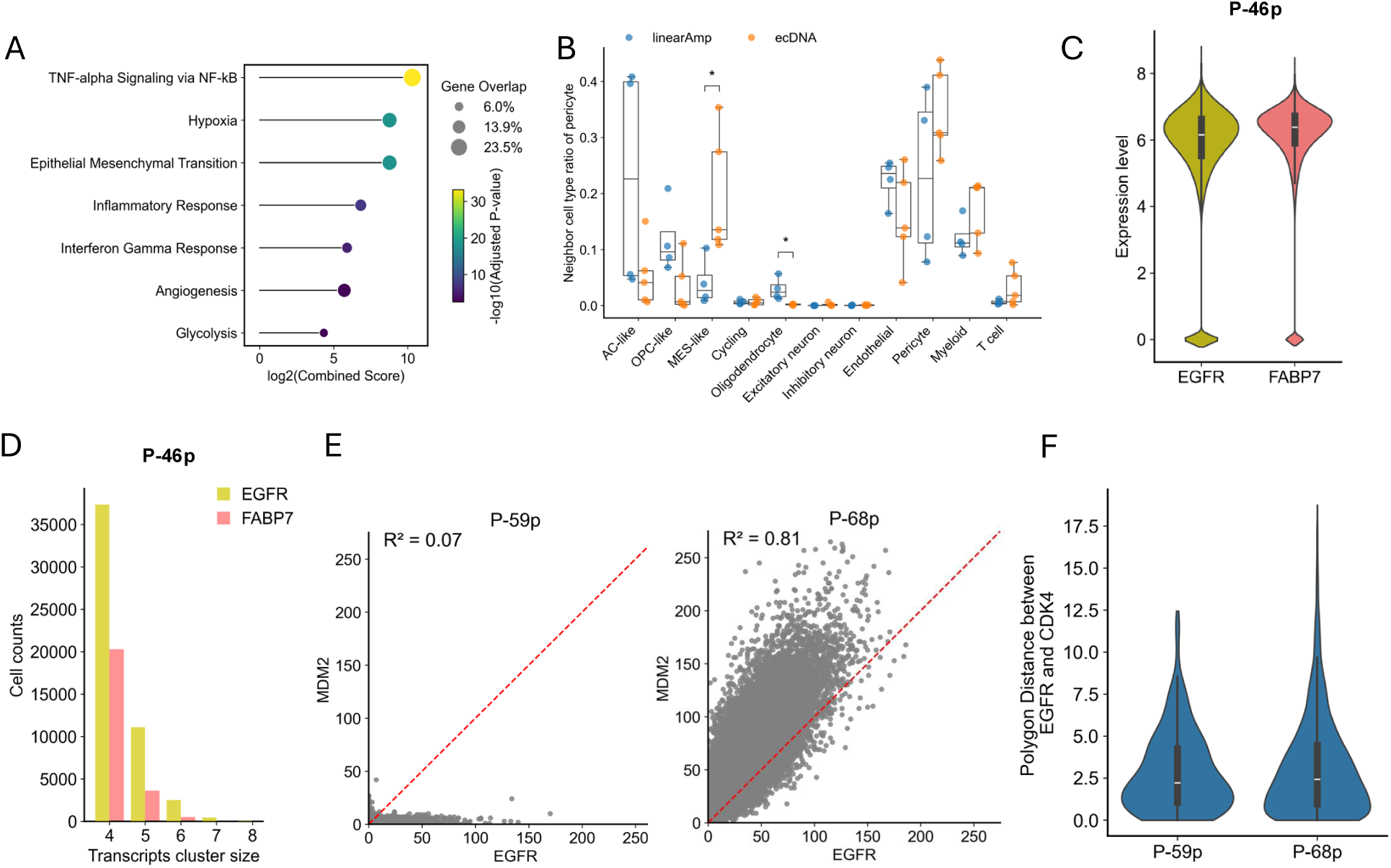
**A)** Bulk RNA-seq data analysis showed that ecDNA-containing tumors have significantly higher hypoxic and metabolic activity compared to linear EGFR-amplified tumors **B)** Cell type composition of tumors with linear versus ecDNA-based *EGFR* amplification shows significant enrichment of MES-like and pericyte cells in ecDNA cases, while AC-like cells predominate in linear cases. P-values were calculated using the two-sided Wilcoxon rank-sum test (* *P* < 0.05). **C)** FABP7 and *EGFR* expression levels are comparable in P-46p tumor Xenium data. **D)** Distribution of *EGFR* transcript cluster sizes across cells reveals larger and more frequent transcriptional clusters than *FABP7* transcription clusters in P-46p tumor Xenium data. **E)** Scatterplots of *EGFR* and *MDM2* expression in tumors P-59p (*EGFR* amplified, *MDM2* not amplified) and P-68p (*EGFR* and *MDM2* co-amplified on same ecDNA) show stronger expression correlation in P-68p. **F)** Distribution of polygon distances between EGFR and CDK4 clusters in P-59p (amplified within separate ecDNAs) and P-68p (amplified within the same ecDNA).

## Methods

### Human Tissue Collection

All patients provided written informed consent under protocols approved by the Institutional Review Board of MD Anderson, in accordance with ethical standards for genomic research. The pathological evaluation confirmed that these tissues were malignant glioma specimen. DNA and RNA extraction was performed using standard methods for fresh frozen tissues. Illumina whole genome sequencing (WGS) (tumor: 60x, normal: 30x), bulk RNA-sequencing was generated by UT MD Anderson Moonshot Program Cancer Genomics Laboratory using standard Illumina library preparations and sequenced on Illumina NovaSeq platform. Oxford Nanopore Technology (ONT) long read sequencing was performed on the PromethION platform by MD Anderson Advanced Genome Sequencing Core. Single nuclei DNA sequencing for 2 pairs of primary and brain metastasis samples was performed by the CPRIT single cell genomic center (SCGC) at MD Anderson Cancer Center. 10x Genomics Xenium spatial transcriptomics were performed with our inhouse machine as described below.

### Short-read WGS Data Processing and Alignment

WGS data pre-processing was conducted in accordance with the GATK Best Practices using Sarek (nf-core-sarek-3.1.2^54^). BWA^55^ (v0.7.17) was used to align the data to the genome (GRCh38/hg38) using bwa mem. For single nucleotide variant (SNV) calling, GATK^56^ MarkDuplicates was used to identify duplicate reads, followed by Quality Score Recalibration (BQSR) using GATK’s BaseRecalibrator and ApplyBQSR to improve base quality scores.

### Fingerprint analysis

Given the multi-omic nature of our study, we implemented a single-nucleotide polymorphism matching algorithm to ensure accurate correspondence between tumor sections and their respective tumors. We performed DNA fingerprint analysis using Picard (v2.27.4) CrosscheckFingerprints^57^. The hg38 haplotype map, a collection of SNP “blocks,” was obtained from the pre-computed fingerprint map repository (https://github.com/naumanjaved/fingerprint_maps). Initially, we conducted cross-checks across all randomly paired WGS samples to identify and correct mismatched samples. Subsequently, we compared fingerprints between WGS, Hi-C and RNA-seq datasets in a pair-wise manner, to confirm that all datasets originated from the same patient.

### Purity and Ploidy Estimation

Purity and ploidy of tumor samples were estimated with ascat^58^ (v4.5.0) using matched tumor and normal samples. An SNP GC correction file was downloaded from the ascat GitHub page. For samples with abnormal ASCAT sunset figures (e.g., p18P, total of three tumors), purity and ploidy were estimated using the mean values from Hatchet^59^, FACETS^60^ (v0.6.2), and Purple^61^ (v3.7.2).

### Somatic Variant Calling

Single nucleotide variants (SNVs) and small insertions/deletions (indels) were identified using GATK Mutect2^56^ (GATK v4.3.0.0) and Strelka2^62^ (v2.9.10) within the Sarek pipeline. To obtain high-confidence SNVs, results from Mutect2 and Strelka2 were merged using pysam. Only SNVs that passed all quality metrics and showed consistent ALT type between the two tools were retained. Variants were annotated with VEP^63^ (v110.0) and SnpEff^64^ (v5.1).

### Structural Variant Calling

Structural variants (SVs) were detected using JaBbA (v1.1), SvABA^65^ (v1.2.0), Purple (v3.7.2), and Smoove (v0.2.5, https://github.com/brentp/smoove). In JaBbA, the coverage ratio was calculated using FragCounter (https://github.com/mskilab/fragCounter), while ploidy and purity were derived from ascatNgs or mean values from ascatNgs, FACETS (v0.6.2) and Purple (v3.7.2) as mentioned in the Purity and Ploidy Estimation section. SV input of JaBbA was from unfiltered results from SvABA. To get high confidence SVs, SVs from these four tools were merged. SVs with the same type and within a 1kb distance at both ends were considered as the same SV. Translocation SVs were only retained if supported by at least three tools, whereas other SVs required support from at least two.

### Driver Mutation Identification

Putative driver mutations in a selected gene list *’IDH1’, ‘IDH2’, ‘EGFR’, ‘CDKN2A’, ‘ATRX’, ‘TERT’, ‘CIC’, ‘FUBP1*’ were identified using GATK Mutect2 (GATK v4.3.0.0) or Purple (v3.7.2). Mutations from Purple with a likelihood greater than 0.75 were retained. Mutect2 results were annotated with VEP, with a maximum distance set to 500 bp.

### ecDNA detection from bulk WGS

To detect ecDNA, all samples in the WGS cohort were analyzed using the AmpliconSuite pipeline (v1.3.4), which integrates AmpliconArchitect (v1.3.r6) and AmpliconClassifier (v1.2.1). The pipeline (*AmpliconSuite-pipeline.py*) was launched using BAM files with default settings. DNA copy number alterations were identified using CNVkit^66^ (v0.9.12), and the resulting CNV calls were used to generate “seed regions.” Seed regions were defined as regions with a copy number ≥4.5, a length >50 kb, and at least 2 copies above the median chromosome arm ploidy, while masking repetitive and poorly mappable sequences. AmpliconArchitect used these seed regions to construct a local genome breakpoint graph, representing genomic copy number and structural variants (SVs). The ultimate amplicon graph is formed by merging regions from other locations joined by additional SVs to this graph’s regions. Paths and cycles were then extracted from the genome graph and analyzed with AmpliconClassifier to predict the presence of ecDNA.

### Bulk RNA-seq analysis

Total RNA was extracted from approximately 20 mg of flash-frozen brain tumor tissue. mRNA was purified from ∼100 µg of DNase-digested total RNA and used to generate sequencing libraries, which were run on the Illumina NextSeq 6000 platform. Processed reads were aligned to the GRCh38 reference genome using STAR (v2.7.9a). Gene-level read counts were calculated using featureCounts (subread v2.0.6). Gene expression levels were quantified as transcripts per million (TPM) based on the read counts.

### Gene enrichment analysis

Differential gene expression analysis was performed using PyDESeq2^67^ (v0.5.0). Genes with |Log2FoldChange| > 1 and padj < 0.05 were considered significantly differentially expressed (DEGs). Enrichment analysis of DEGs was conducted with GSEApy.enrichr^68^ (v1.1.7) using the Hallmark pathways database from MSigDB. Additionally, ssGSEA was performed with GSEApy.ssgsea, and the NES scores for the top Hallmark pathways in each sample were obtained.

### Hi-C Data Processing and Alignment

Hi-C reads were aligned to the human reference genome (GRCh38/hg38) using BWA (v0.7.17) with bwa mem with the -SP5M flag. To generate Hi-C .cool files, the BAM files were first converted to a binned contact matrx files using Pairtools. These matrix files were then converted to.mcool files using Cooler^69^. Resolutions of 5k, 10k, 50k, and 100k were retained. Two different normalizations were applied: the first using the ICE (Iterative Correction and Eigenvector) algorithm with cooler balance, and the second using CNV balance by NeoLoopFinder’s correct-cnv.

### Hi-C Structural Variant Calling and Neoloop Identification

SVs were identified from the Hi-C ICE-balanced .cool files using Eagle-C^70^ by analyzing changes in contact patterns between genomic regions. The parameters for Eagle-C were set to --prob-cutoff-5k 0.8, --prob-cutoff-10k 0.8, and --prob-cutoff-50k 0.99999. Neoloops were identified using neoloop-caller with --prob set to 0.99 and --no-clustering.

### Single-nucleus whole-genome sequencing

Single nucleus whole genome sequencing (snWGS) was generated from frozen tissue samples with the ARC-well protocol. Data was aligned to hg38 with BWA mem 2. Samtools sorting, picard duplicate marking, and samtools indexing were performed. ASCAT.sc (v0.1) was used to call copy number alterations with the multiPCF setting for consistent segmentation between cells. Ploidies were fit accordingly to their DAPI FACS peak. Peaks that resembled diploid or G2/M were fit to a ploidy range of 1.5-2.5. Peaks that resembled a whole genome duplication were fit to a ploidy range of 3.5-6. Peaks that weren’t clearly distinguishable between G2/M and whole genome duplications were fitted to a broader range of 1.5-6 to account for the uncertainty. The focal amplification region was specified with the svinput argument, which contained the largest interval from AmpliconArchitect cycles file. A modified version of the ASCAT.sc quality control function was used to select high quality cells for further analysis with the following settings: at least 2^16 reads per cell, the upper 90% of cell qualities, and within 0.03 distance of a line fitting the cells between number of reads and logR noise. Copy number segments were adjusted to be roughly proportional to their genomic length for plotting, which was performed with ComplexHeatmap^71^. In order simplify the main figure for visualization, further filtering was performed to remove cells that were nearly diploid (healthy cells) or highly aneuploid (like whole genome duplicated cells). To do this, cells with less than 25% or more than 75% of their genomic segments equal to 2 were filtered out. All cells that passed quality control are visualized in the supplementary figures.

### Single-nucleus Multiome

Single-nucleus Multiome (ATAC + Expression) analysis was performed on tumor samples. Nuclei were isolated from approximately 30 mg of freshly frozen tissue using the 10x Genomics Nuclei Preparation Kit, with slight modifications. Briefly, tissue samples were thawed in 200 µL of the nuclei lysis buffer, cut into small pieces, and homogenized with a pestle until no visible tissue fragments remained. The samples were then incubated on ice for 5–10 minutes with an additional 300 µL of nuclei lysis buffer. After centrifugation, nuclei were washed twice with wash buffer, resuspended in diluted nuclei buffer, and counted. Library preparation followed the 10x Genomics user guide (CG000338), and sequencing was performed on the Illumina NovaSeq 6000 platform. Reads were aligned to the human genome (GRCh38) using cellranger-arc (v2.0.0, 10x Genomics) to generate UMI counts and ATAC-seq fragments. snRNA cells were filtered as: 30000 ≥ UMI counts ≥ 500, 6500 ≥detected gens ≥ 200, 10 > Mitochondrial count percentage, doublet removal (predicted by Scrublet). Genes detected in less than 3 nuclei were also removed. snATAC cells were filtered as: 100000 ≥ counts ≥ 1000, TSSe ≥ 5 and doublet removal (predicted by Scrublet). Features in the blacklist (hg38-blacklist.v2) were excluded. Finally, only cells present in both the snRNA and snATAC datasets were retained for downstream analysis. snRNA data were analyzed using Scanpy^72^ (v1.10.2), while snATAC data were processed with SnapATAC2^73^ (v2.7.0). For snRNA data, UMI counts were normalized to a target sum of 10,000 and log transformed. The 2,000 most variable genes were selected for principal component analysis (PCA). Data integration was performed using harmonypy (v0.0.10), followed by clustering with the Leiden algorithm. Marker genes for each cluster were identified using *scanpy.tl.rank_genes_groups* with the default Wilcoxon method. Cells were manually annotated based on marker gene expression, and these annotations were also mapped to the snATAC dataset. For snATAC data, the cell × feature count matrix was reduced in dimensionality using *snapatac2.tl.spectral*, integrated with *snapatac2.pp.harmony*, and clustered using *snapatac2.tl.leiden*. Copy number variations (CNVs) were inferred from snRNA data using infercnvpy (v0.5.0) and from snATAC data using epiAneufinder^74^ (v1.0.1).

### Oxford NanoPore Long-read Sequencing

Oxford Nanopore Technology (ONT) sequencing was performed with the R10.4 flowcell. Basecalling and alignment to the hg38 genome was performed using Dorado (v0.9). The aligned bam was sorted, MD tags were added, and indexed with Samtools. Bams were merged into a single bam file for each sample. Nanoplot^75^ was used to obtain sequencing quality information. Coral was used to reconstruct ecDNA amplifications with default settings with two adjusted parameters: coral seed --gain 10, coral reconstruct --min-bp-support 5, 10. Both 5 and 10 were used for --min-bp-support because they resulted in some samples with distinct AmpliconClassifier results. Coral reconstruct --min-bp-support 10 results were used further analysis. AmpliconClassifier were then used to classify focal amplifications.

### Spatial transcriptomics with 10x Xenium Analyzer

Fresh Frozen (FF) and Formalin-Fixed Paraffin-Embedded (FFPE) tissues were used in this study. FF tissues embedded in OCT were sectioned to a thickness of 10 µm onto Xenium slides and processed according to the 10x Genomics user guide (CG000581) for fixation and permeabilization. FFPE blocks were sectioned to a thickness of 5 µm, placed on Xenium slides, and processed following the user guide (CG000580) for deparaffinization and decrosslinking.

Xenium in situ gene expression analysis was performed according to the appropriate 10x Genomics user guides, either without cell segmentation staining (CG000582) or with cell segmentation staining (CG000749). A custom gene panel of 100 genes (Supplemental Table 5) was added to the pre-designed human brain gene panel. A mix of pre-designed and custom gene expression probe hybridization was conducted at 50 °C overnight. Unhybridized probes were removed through multiple washing steps, after which the probes were ligated and Rolling circle amplification (RCA) primers were annealed at 37 °C for 2 hours to generate circularized padlock probes, which were subsequently amplified at 30 °C for 2 hours. If cell segmentation staining was used, it was performed at 4 °C overnight according to the CG000749 user guide. Lastly, background fluorescence was quenched, nuclei were stained, and the prepared slides were loaded onto the Xenium Analyzer. The Xenium Analyzer performed multiple rounds of fluorescence probe hybridization, imaging, and probe removal automatically. Z-stacked fluorescence images were acquired and analyzed using Xenium Onboard Analysis (v3.0.0), which included decoding, cell segmentation, analysis, and quality control (QC).

After the Xenium run, slides were stored hydrated in PBS-T (0.05% Tween-20 in PBS) at 4 °C. For H&E staining, slides were processed according to the user guide (CG000613), mounted with coverslips, dried, and scanned with a slide scanner (PE Vectra Polaris) at 20x magnification.

### DNA-FISH on post-Xenium slides

Post-Xenium FFPE slides were treated with Quencher Removal solution as suggested by 10x Genomics prior to performing Fluorescence in situ hybridization (FISH). Slides were processed according to the Agilent FISH Protocol on FFPE Samples using the DAKO Histology FISH kit. A set of commercially available probes from MetaSystems and Leica Biosystems were used to target and hybridize with specific genes to confirm abnormalities observed previously by Xenium and molecular analysis. Centromere probes, labeled in green, were used as controls, while locus-specific probes targeting genes were labeled in red. Cell and signal imaging were conducted using an ECHO REVOLVE R4 fluorescent microscope.

### Xenium data processing

We performed Xenium spatial transcriptomics on 62 samples using either probes combined with the human brain base probe panel. Xenium Analyzer output data (transcript locations, gene assignments, cell x gene tables, cell/nucleus boundaries, and stacked nuclei images) were processed using the SpatialData framework^76^ (v0.2.2). Individual samples within each Xenium dataset were separated using spatialdata.polygon_query and stored in Zarr format.

For datasets using P1 probes without cell segmentation staining, the cell segmentation was initially performed by expanding nuclear boundaries by 5 μm in XOA. To optimize cell segmentation, we compared two re-segmentation methods: Baysor (v0.7.0) and Proseg (v2.0.2). Proseg produced superior clustering results, so Proseg generated cell x gene tables were used for downstream analysis. Cells with ≤20 or ≥800 transcripts, or ≤4 detected genes were filtered out. Leiden clustering was then applied to each sample using Scanpy (v1.10.2) after normalizing to a target sum of 10,000 and log transformation. Clusters with median transcript counts ≤50 were excluded. The cell x gene tables from all samples were merged into an AnnData object (v0.10.8), normalized (target sum = 10,000) and log transformed. Gene Expression Program (GEP) analysis was conducted using cNMF^77^ (v1.5.4) with Harmony batch correction. cNMF was run for 500 iterations with k values from 5 to 15, with k = 11 selected as the optimal GEP clustering number. The top 15 marker genes from each GEP were selected and duplicates removed, resulting in 155 unique marker genes. A GEP was assigned to a cell if it had minimum 10% usage and higher usage than any other program. These GEP assignments were used for cell type annotation. For FFPE tissue p46P, which used P1 probe with cell segmentation staining, GEP usage derived from cNMF analysis was transferred using starCAT^78^ (v1.0.9). The GEP with the highest program usage determined the cell type annotation.

Main marker gene’s used for cell annotations include: AC-like *(EGFR, FABP7, MT3, GFAP, AQP4, TTYH1);* OPC-like *(BCAN, OLIG1, OLIG2, PDGFRA, SOX8, SOX4, SOX11); MES-like (IGFBP5, HILPDA, VEGFA, NDRG1);* Oligodendrocyte *(MAG, CNDP1, SOX10, MOG, CNTN2, ERMN, MOBP, MAL, CLDN1);* Neuron *(SNAP25, ANO3, SLC17A7, C1QL3, GAD1, GAD2, ANK1);* Myeloid *(TMEM119, P2RY12, CX3CR1, CD163, MSR1, MRC1, C1QA, C1QB, IL1B);* Cycling *(CCNB2, CDK1, CENPF, MKI67, TOP2A);* Endothelial *(PECAM1, FLT1, PHLDB2, CLDN5);* Pericyte *(COL12A1, COL3A1, PDGFRB, DCN, FBLN1);* T cell *(CD3D, CD8A, CD8B, FOXP3, CCL5, CD2, CD3G, CXCR4, CYTIP, CD48, GZMA, TRAC*).

### Neighborhood analysis

Spatial niche analysis was conducted using CellCharter^79^ (v0.3.1) and Banksy^80^ algorithms. To identify the optimal niche number for each sample, CellCharter encoded cells into a network where nodes represented cells and edges represented cell connectivity. Neighborhoods for each cell were aggregated into a single feature to the layer of 4 using the gr.aggregate_neighbors function. A Gaussian Mixture Model was created and fitted to predict niche labels for all cells, with the optimal niche number determined by tl.ClusterAutoK. Banksy was then run with the identified niche number and a lambda value of 0.8 to define spatial niches. To identify consistent niches across all sample cohorts, we computed cell type composition ratio within each niche, performed Pearson correlation analysis between niches, and applied hierarchical clustering. This analysis revealed eight distinct meta-neighborhoods characterized by high intra-group correlation.

We assessed spatial proximity between cell types on the connectivity graph using CellCharter’s gr.nhood_enrichment function. The analysis focused on malignant (AC-like, OPC-like, and MES-like) cells and two subtypes of myeloid cells (inflammatory microglia and immunosuppressive scavenger). The normalized enrichment score for each cell type pair was calculated within individual samples, and the scores were averaged across samples for final evaluation.

### Spatial coherence analysis

The spatial coherence score, a measure of the structural organization or disorganization within a tumor, was defined previously^22^. To calculate the score, the tissue sample with cell annotations was evenly divided into 200 x 200 bins, and the number of cells from each cell type was determined for each bin. The dominant cell type within each bin, defined as the one with the highest cell count, was assigned as the program for that bin. The coherence score was calculated by examining each bin and its surrounding 3 x 3 square neighbors. If the neighboring bins shared the same program as the central bin, the bin was assigned a score of 1; otherwise, it was assigned a score of 0. This process was repeated for all bins and programs. The total scores for each program were summed and divided by the total number of bins associated with that program. The resulting ratios were scaled between 0 and 1, and the average of these scaled ratios was used as the overall coherence score for the sample.

### Neighboring cells analysis

In each sample, the 8 nearest neighboring cells of each pericyte cell were identified using KDTree (scipy, v1.14.1). Neighboring cells located more than 200 µm away were excluded. The remaining neighboring cells were aggregated to calculate the ratio of different cell types. The nearest AC-like or MES-like cell to each pericyte was identified within a 200 µm radius, and their Euclidean distance was measured.

### Gene transcripts clustering analysis

The X and Y coordinates (2D) were used to analyze gene transcript clustering within each cell. For each transcript, all other transcripts of the same gene within 1 µm were identified and counted. A transcript was classified as part of a cluster if it had at least 3 neighboring transcripts. The frequency of clusters of all sizes was then recorded. A polygon representing each cluster was generated using Shapely (v2.0.4), and the distance between EGFR and CDK4 polygons, or between EGFR and nucleus polygons, was calculated for each cell.

### Patient-derived glioma stem cell lines

Following thawing at 37°C, the cells were resuspended and cultured in DMEM/F12 media supplemented with B-27, hEGF, hbFGF, and antibiotics (McDonald, M. F. et al., 2024). The cultures were maintained in an incubator at 37°C with 5% CO2. Once the cells reached approximately 80% confluency, they were passaged into new flasks. For harvesting, the cells were treated with colcemid and exposed to a hypotonic KCl solution. Carnoy’s fixative solution (3:1 methanol:acetic acid) was added, and the cell pellets were stored at 4°C until slide preparation.

### Fluorescence in situ Hybridization (FISH)

Slides were labeled with appropriate identification details, and two drops of the cell suspension were applied. The slides were then dried on a slide warmer before being aged overnight at 60°C. Following this, the slides were processed for FISH. A set of commercially available probes including *EGFR*(7q11.2)/SE7 (D7Z1-centromeric region), RUO-*MDM2*(12q15)/SE12(D12Z3), RUO-MYCN/SE2 from Leica Biosystems were used to study the mutations in each cell line. Centromere probes, labeled in green, were used as controls, while locus-specific probes were labeled in red, except for PAX6, which was also labeled in green. The signal analysis was performed using an ECHO REVOLVE R4 fluorescent microscope and 100 interphase cells were analyzed for each FISH probe.

